# Structure and Activity of Class II Lanthipeptides from a Thermophilic Bacterium

**DOI:** 10.64898/2026.04.02.716199

**Authors:** Enleyona Weir, Lingyang Zhu, Wilfred A. van der Donk

## Abstract

Lanthipeptides represent a large group of ribosomally synthesized and post-translationally modified peptides (RiPPs). They offer promising avenues for discovering new antibacterial and antifungal agents. Here, we identify and structurally analyze the product of the *tla* BGC, which encodes a class II lanthipeptide in the thermophilic bacterium *Thermoactinomyces* sp. DSM 45891. Co-expression of the lanthipeptide synthetase TlaM with its substrates in *Escherichia coli* resulted in modification of the two precursor peptides TlaA1 and TlaA2, which share 58% sequence identity. TlaA1 was dehydrated up to seven times, with the major products having undergone five and six dehydrations, whereas TlaA2 was dehydrated seven times. In both peptides, four thioether rings were formed with two overlapping DL-(methyl)lanthionine rings at the C-terminus. Both peptides also contain two non-overlapping DL-methyllanthionines near the N-terminus and in the center of the peptide. These peptides deviate from the general rule of stereoselective LL-(methyl)lanthionine formation from a Dhx-Dhx-Xxx-Xxx-Cys motif (Dhx = dehydroalanine or dehydrobutyrine). AspN-cleaved TlaM-modified TlaA1 displayed anti-microbial activity against a subset of bacteria including Gram-negative ESKAPE pathogens. We named the lantibiotic thermolanthin.

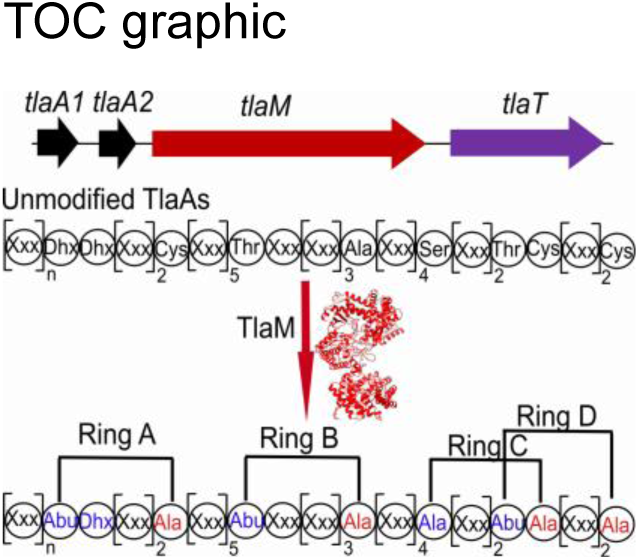

## 1. Introduction

Lanthipeptides represent a large group of ribosomally synthesized and post-translationally modified peptides (RiPPs) characterized by intramolecular thioether crosslinks called lanthionine and methyllanthionine.[^1, 2^] Their wide-ranging functions, including antimicrobial and antifungal activity, virulence functions, and signaling roles,[^3^] and their promise for bioengineering,[^4–7^] have motivated researchers to discover new members by genome mining exercises.[^8–10^] Lanthipeptides with antibacterial activity have been termed lantibiotics.[^11^] Lanthipeptide biosynthetic enzymes play a key role by introducing specific ring patterns that impart the various activities.[^12^] A previous large-scale bioinformatic genome mining effort focused on lanthipeptide biosynthetic gene clusters (BGCs) and revealed high sequence diversity amongst precursor peptides across class I-IV lanthipeptides.[^13^] The post-translational modifications (PTMs) take place in the C-terminal segment of the precursor peptide called the core peptide. An N-terminal leader peptide is removed after completion of the PTMs to release the final mature lanthipeptide.[^14^]

In most reported lanthipeptide BGCs, a single precursor peptide is encoded that is morphed into the mature natural product. For a relatively small subset of the BGCs, two precursor peptide genes are encoded within a single BGC.[^13^] These typically produce two-component lanthipeptides that act synergistically to achieve antimicrobial activity.[^15, 16^] For such two-component systems, the two peptides usually have considerable sequence diversity and result in two products with different ring patterns. The individual peptides typically have little or no activity in the absence of their partner peptides. For instance, the two-component lantibiotics lacticin 3147 (Ltnα/Ltnβ), lichenicidin (Licα/Licβ), haloduracin (Halα/Halβ), cytolysin (CylL_L_”/CylL_S_”), plantaricin W (Plwα/Plwβ), and roseocin (Rosα/Rosβ) require both distinct post-translationally modified peptides for physiological activity.[^17–28^] The individual peptides of such two-component systems can be produced either by a single enzyme, capable of modifying both peptides,[^29–31^] or by two distinct enzymes, each modifying one of the two precursor peptides.[^21, 24, 32^]

In other lanthipeptide BGCs, multiple precursor peptides are encoded that are identical or highly similar in sequence that result in a group of congeners with the same ring patterns.[^13^] It is likely that these closely related products expand the spectrum of activity or, in the case of identical precursor sequences, increase productivity by increasing the ratio of precursor peptides compared to biosynthetic machinery without placing the two under different promoters and create a specific operon structure that complicates regulation. An alternative solution to increase diversity and/or productivity in the RiPP field has been the evolution of precursor peptides with multiple core peptides,[^33, 34^] but this strategy has not yet been reported for lanthipeptides. Finally, examples have been reported, especially in cyanobacteria, where multiple precursor peptides or multiple core peptides are encoded in the BGC to generate products with independent activities that greatly increase the structural and functional diversity of the output of RiPP BGCs.[^35–37^]

Here we describe and characterize TlaM, a class II bifunctional lanthipeptide synthetase from the thermophile *Thermoactinomyces sp.* DSM 45891 that performs both dehydrations and cyclizations in TlaA1 and TlaA2 precursor peptides encoded in the *tla* BGC. The two precursor peptides share 57.9% sequence identity, which is relatively low for BGCs that encode a set of congeners of the same lanthipeptide and suggesting that possibly this BGC could produce a two-component lantibiotic. TlaM is a member of the class II lanthipeptide synthetases that use an N-terminal domain to phosphorylate Ser and Thr residues and subsequently to eliminate the phosphate to generate dehydroalanine (Dha) and dehydrobutyrine (Dhb) residues, respectively.[^38, 39^] The C-terminal domain contains a Zn^2+^ in its active site that is important for catalyzing Michael-type additions of select Cys residues to the Dha and Dhb residues to form thioether crosslinks called lanthionine (Lan) and methyllanthionine (MeLan), respectively.[^40^]

Thermophilic bacteria represent a promising source of robust, adaptable biosynthetic machinery. Enzymes from thermophiles often exhibit elevated stability, enhanced tolerance to structural variation, and favorable biochemical resilience given their ability to function at elevated temperatures.[^41–45^] Only a few examples of lanthipeptides have been reported that are produced by thermophilic bacteria.[^46^] One such example, geobacillin I, is a class I lanthipeptide produced by *Geobacillus thermodenitrificans* NG80-2, which contains seven (methyl)lanthionines. Geobacillin I exhibits antimicrobial activity comparable to that of the commercial food preservative nisin A, but with greater thermal and pH stability.[^47^] Geobacillin II, a class II lanthipeptide from the same organism that undergoes modification by the bifunctional enzyme GeoM, has a different ring topology than geobacillin I.[^47^] In this study, we used *Escherichia coli* to heterologously produce the two lanthipeptides from the *tla* BGC. We demonstrate that TlaM-modified TlaA1 and TlaA2 contain (methyl)lanthionine crosslinks and dehydroamino acid residues. In addition, the ring patterns of the products were characterized by nuclear magnetic resonance (NMR) spectroscopy and mass spectrometry, and their biological activity after leader peptide removal was assessed. These lanthipeptides do not have structural homologs among known lanthipeptides.

## 2. Results and Discussion

### 2.1 Bioinformatic identification of the *tla* biosynthetic gene cluster

The *tla* BGC was chosen from a previously reported sequence similarity network of class II lanthipeptides with the expectation that it could be the first characterized two-component lanthipeptide from a thermophilic organism.[^13^] Using the RODEO webtool 2.0,[^48^] two precursor peptides, encoded by *tlaA1* and *tlaA2*, were predicted to consist of 69 and 74 amino acid residues, respectively (**Figure 1**). No additional modifying enzymes were encoded near these peptides with the exception of a canonical class II lanthionine synthase, encoded by *tlaM* and a peptidase-containing ATP-binding cassette transporter (PCAT)[^49^] ThtT predicted to remove the leader peptide from the modified precursor at a double Gly motif using its N-terminal C39 peptidase domain.[^50^]

**Figure 1.**
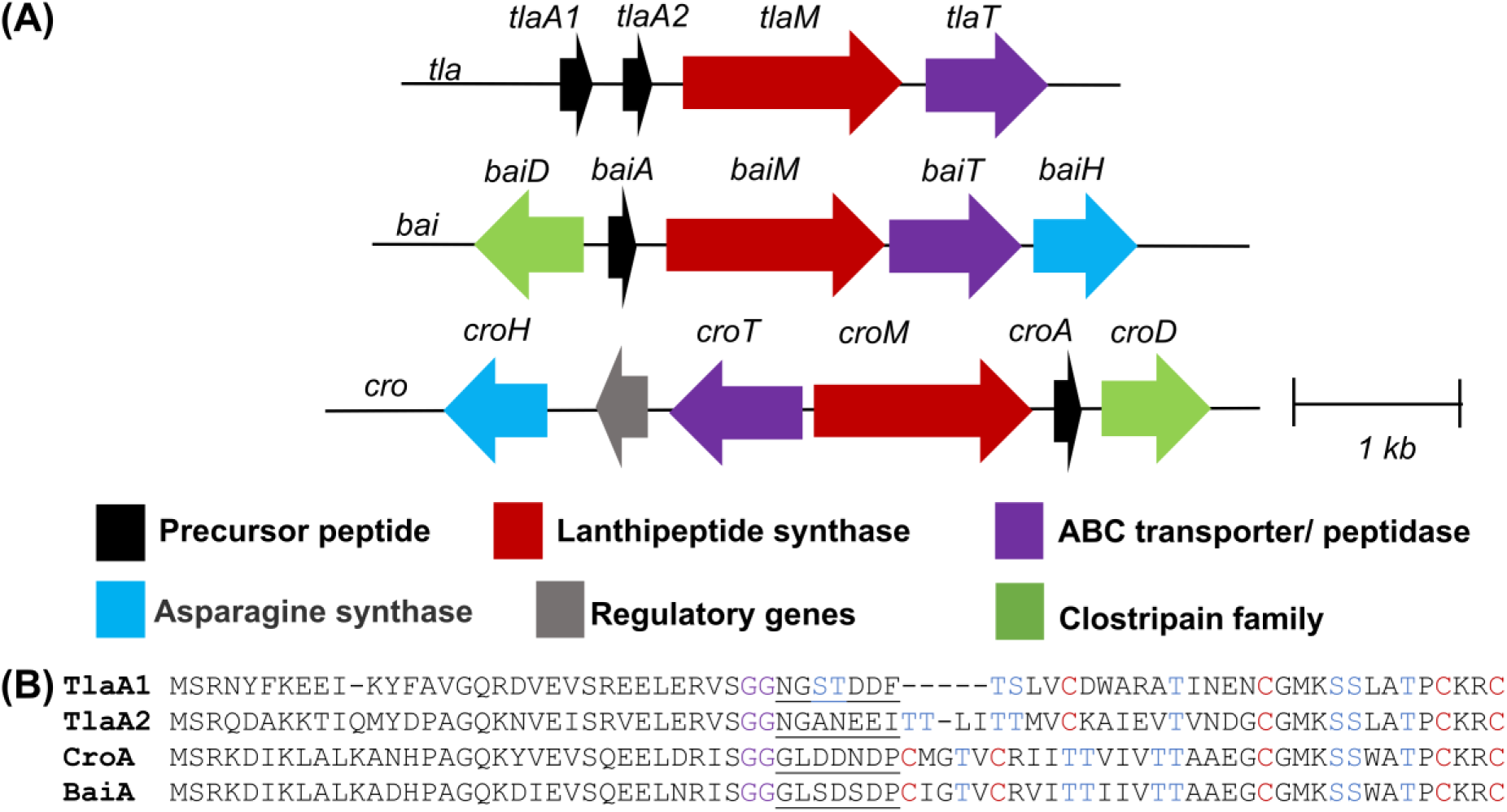
The *tla* biosynthetic gene cluster from *Thermoactinomyces* sp. DSM 45891 and other orthologous BGCs (accession numbers in **Table S1**). A) Gene composition of the *tla*, *bai,* and *cro* BGCs. The genes encoding the precursor peptides are in black, LanMs in red, and PCATs in purple. B) Precursor peptide sequences from each BGC; Ser/Thr and Cys residues, shown in blue and red, respectively, were expected to be involved in post-translational modifications. Double glycine motifs are shown in purple. The amino acid residues after the double Gly cleavage site and before the first Cys or dehydrated amino acid are underlined and may be involved in a second proteolytic cleavage event.

TlaA1 and TlaA2 both contain eight Ser/Thr amino acid residues and four Cys residues in the predicted core peptides (**Figure 1B**). The constellation of these residues requires a product distinct from any previously characterized lanthipeptide. A BLAST search retrieved very few peptides with sequence homology to TlaA1 and TlaA2. Two putative precursor peptides BaiA and CroA from *Baia soyae* and *Croceifilum oryzae,* respectively, display sequence homology towards their C-termini but not at their N-termini (**Figure 1B**). Their BGCs differ from the *tla* BGC, as they encode additional genes, including an asparagine synthase and a clostripain C11 peptidase. The former may form lactam,[^51^] amide[^52, 53^] or nitrile[^54^] structures based on previous roles in RiPP biosynthesis, and the latter may be responsible for a second proteolytic cleavage event after PCAT processing to produce the mature product of the *bai* and *cro* cluster.[^14^] Such second proteolytic cleavage events are common for two-component lantibiotics and typically involve the removal of six or seven amino acids. The TlaA1/A2, CroA, and BaiA peptides all contain a heptapeptide between the predicted double Gly cleavage site for a PCAT enzyme and the first Ser/Thr/Cys residue that is likely modified during maturation (**Figure 1B**). The presence of a second protease in the *cro* and *bai* BGCs provides indirect support for a step-wise multiproteolytic process. A sequence alignment of the lanthionine synthetases in the homologous BGCs showed relatively similar sequences. TlaM and BaiM share 62.4% identity, whereas TlaM and CroM have 62.6% identity (**Figure S1**). Hence, these lanthipeptide synthetases are hypothesized to produce similar ring patterns arising from their homologous C-terminal substrate sequences.

### 2.2 Heterologous production of modified TlaA1 and TlaA2 in *E. coli*

*E. coli* was used as heterologous host for co-expression of *tlaA1* and *tlaA2* with *tlaM*. The genes encoding TlaA1 and TlaA2 were separately inserted into pET-Duet vectors. The precursor peptides were fused at their N-termini to a histidine tag for purification and a small ubiquitin-like modifier (SUMO) tag to improve yield[^55^] to produce His_6_-SUMO-TlaA1 and His_6_-SUMO-TlaA2. The lanthipeptide synthetase gene *tlaM* was cloned into multiple cloning site (MCS) 1 of a pRSF-Duet-1 vector, resulting in an N-terminal hexa-histidine-tag. After heterologous expression of His_6_-SUMO-TlaA1 and purification by immobilized metal affinity chromatography (IMAC), both the His_6_-SUMO and the putative leader peptide were removed *in vitro* with the purified C39 protease LahT150,[^56, 57^] an ortholog of the protease domain of TlaT. The expected peptide corresponding to the core peptide sequence was observed by matrix-assisted laser desorption/ionization time-of-flight mass spectrometry (MALDI-TOF MS) (**Figure 2A)**. Coexpression of TlaM and SUMO-TlaA1 followed by LahT digestion resulted in five, six, and seven dehydrations as determined by MALDI-TOF MS (**Figure 2A**). Extracted ion chromatograms (EICs) of LahT-digested TlaM-modified TlaA1 revealed peak area ratios of 1:10:10 for the 7-, 6-, and 5-fold dehydrated products, respectively (**Figure S2A**). Similarly, we co-expressed TlaM and His_6_-SUMO-TlaA2, followed by cleavage with LahT150, and observed seven dehydrations as the major product (**Figure 2B**). EIC analysis confirmed that the seven-dehydrated peptide was the major product formed (**Figure S2B**).

**Figure 2.**
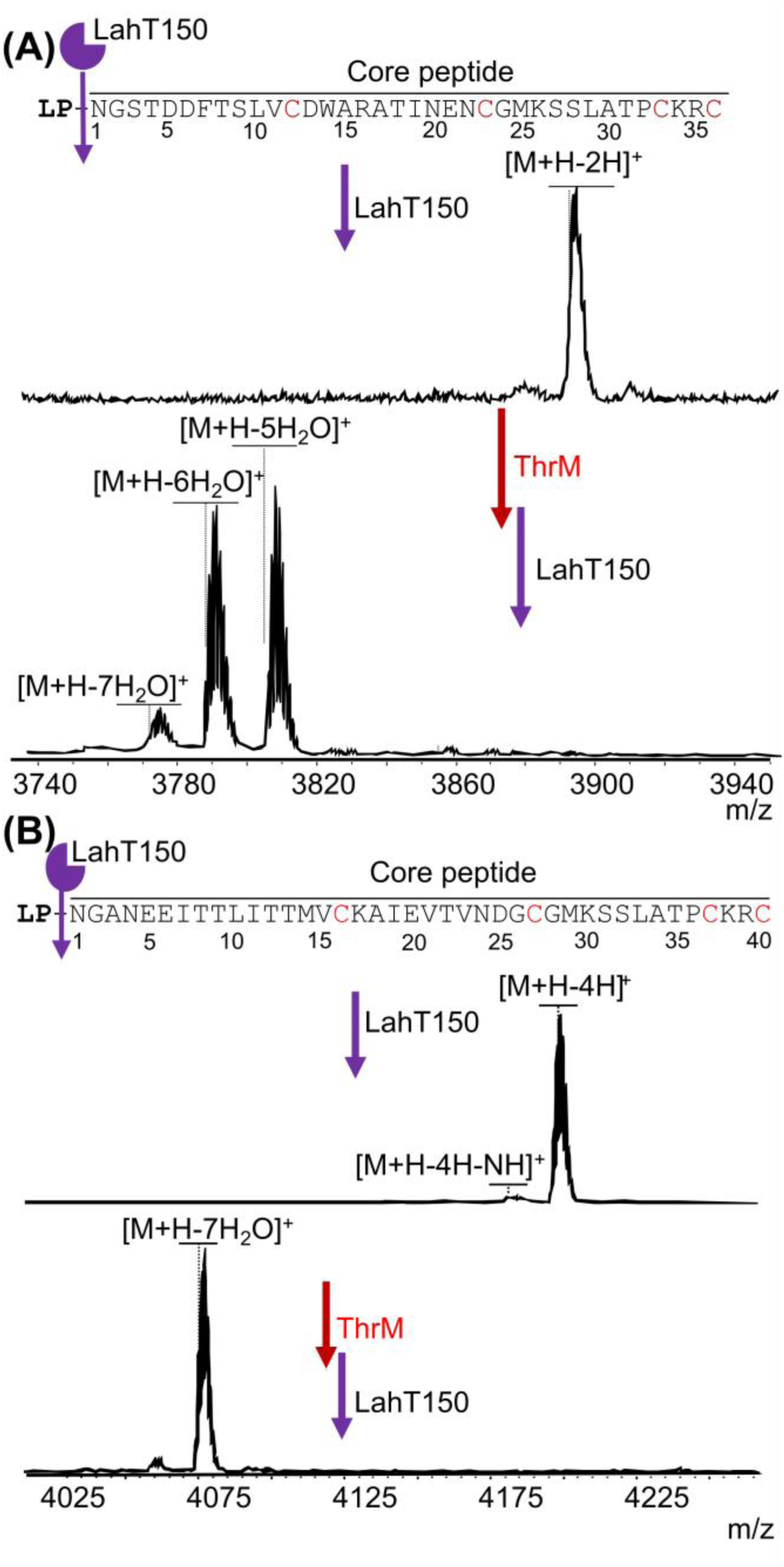
MALDI-TOF mass spectra of the precursor peptides in the *tla* BGC with and without co-expression with TlaM in *E. coli*, followed by *in vitro* cleavage with LahT150 to remove the leader peptide (LP). A) LahT-cleaved TlaA1, observed [M+H]^+^ m/z 3891.2; calculated 3894.7. The difference between calculated and observed values is due to partial formation of disulfide bonds from the four cysteine residues. After heterologous expression with TlaM and LahT cleavage, a mixture of dehydrations was observed. observed [M+H−5 H_2_O]^+^ m/z 3804.9; calculated 3804.7; [M+H−6 H_2_O]^+^ m/z 3787.9; calculated 3786.7; [M+H−7 H_2_O]^+^ m/z 3770.9; calculated 3668.7. B) MALDI-TOF MS of LahT-cleaved TlaA2 (observed [M+H]^+^ m/z 4196.9; calculated 4202.0) and after heterologous expression with TlaM, and LahT cleavage (observed [M+H−7 H_2_O]^+^ m/z 4075.1; calculated 4075.9).

### 2.3 NEM and DTT assays

To determine whether all four Cys residues in TlaA1 and TlaA2 were cyclized and to assess whether any dehydro amino acids were formed after TlaM modification, the LahT150-cleaved TlaA products were subjected to chemical derivatization using *N*-ethylmaleimide (NEM) and dithiothreitol (DTT).[^1^] For TlaA1 treatment with NEM did not result in the formation of an adduct, indicating the peptide was fully cyclized with all four Cys residues forming thioethers with four dehydroamino acids. The modified product was then subjected to a DTT assay.

MALDI-TOF MS analysis demonstrated the presence of a mixture of products with one and two DTT adducts (**Figure 3A**). These observations were consistent with the five and six dehydrations and four cyclizations determined from the NEM assay. To determine the ring pattern and presence of Dha/Dhb residues, the modified TlaA1 was subjected to proteolytic cleavage. TlaM-modified SUMO-TlaA1 (which we will hereafter refer to as mTlaA1) was treated with chymotrypsin, and two fragments were observed that encompass the core peptide (**Figure 3B**). Two and three dehydrations were observed in fragment **1,** whereas fragment **2** was dehydrated three times, showing that the heterogeneity observed in the full length products (mostly 5- and 6-fold dehydration) arises from the N-terminal fragment **1**. Based on the DTT and NEM assays described above, fragment **1** contains one ring that we term ring A, and fragment **2** contains three rings that we term rings B, C, and D. To determine the location of dehydration, mTlaA1 was also digested with endoproteinase GluC, followed by LysC resulting in two fragments A and B (**Figure S3**). Tandem MS analysis of fragment A demonstrated dehydration of Thr4, Thr8, Ser9, and Thr18, with a minor amount of dehydration of Ser3 (**Figure S4**). Thus, the five- and six-fold dehydrated peptides in Figure 2A differ in whether Ser3 is dehydrated or not. The C-terminal fragment B underwent two dehydrations, but their locations could not be determined because of a lack of fragmentation (**Figure S5**), which usually is the result of overlapping rings.[^58^] The two dehydrations in fragment B would originate from Ser27, Ser28, or Thr31.

**Figure 3.**
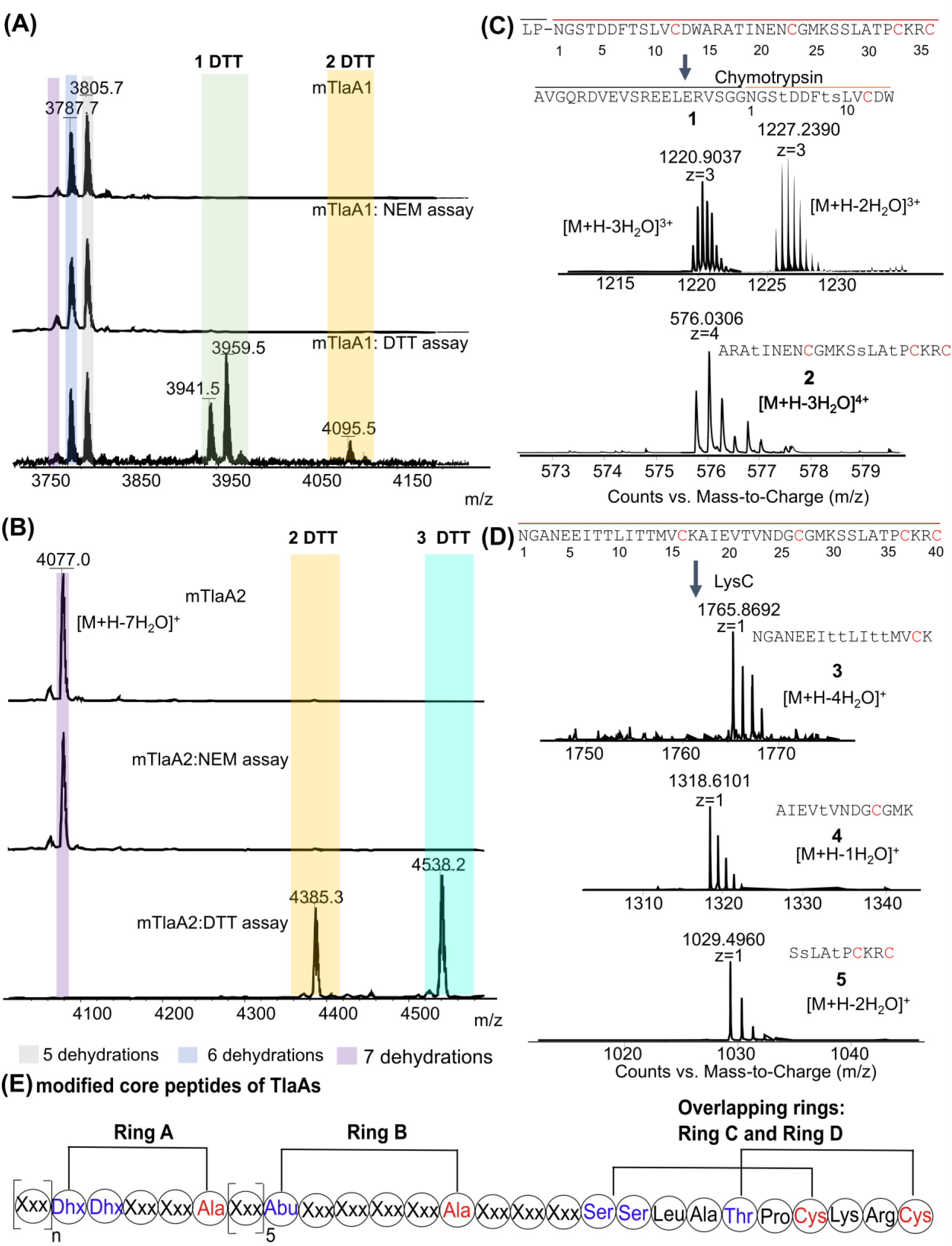
Mass spectrometric analysis of mTlaA1 and mTlaA2 after chemical modification and proteolytic cleavage. A) MALDI-TOF mass spectra of NEM and DTT assays of mTlaA1 cleaved with LahT150. One (+154 Da) and two DTT adducts (+308 Da) were observed. B) MALDI-TOF mass spectra of NEM and DTT assays of mTlaA2 cleaved by LahT150. C) Proteolytic cleavage of mTlaA1 using chymotrypsin resulted in two fragments, **1** and **2**. D) Proteolytic cleavage of mTlaA2 using endoproteinase LysC resulted in three fragments, **3, 4,** and **5**. Lowercase letters represent the identified dehydrated amino acids. E) Schematic representation illustrating the proposed ring patterns of the modified LahT-cleaved TlaAs based on tandem MS data. NMR characterization unambiguously determined the ring patterns as described later.

mTlaA2 cleaved with LahT150 was also subjected to both NEM and DTT assays followed by analysis by MALDI-TOF MS. Adducts were not observed in the NEM assay, indicating that all four cysteines were cyclized. Two and three DTT adducts were observed, suggesting that three dehydroamino acid residues are not involved in cyclization (**Figure 3C**). One of the dehydro amino acids is likely a Dhb, which reacts sluggishly with DTT, especially when located within a (methyl)lanthionine ring, thus explaining incomplete labelling by DTT.

Proteolytic cleavage with AspN and subsequent tandem MS analysis demonstrated that Thr8 and Thr9 were dehydrated in mTlaA2 and are not involved in cyclization (**Figure S6**). Additionally, Thr12 and Thr13 are dehydrated and likely part of a MeLan given the lack of fragmentation between former Thr12 and Cys16 (**Figure S6**). Furthermore, Thr22 is likely dehydrated and part of a ring with Cys27.

As observed for fragment B of mThrA1, the lack of fragmentation in the C-terminal segment of mThrA2 precluded definitive assignment of the final two dehydrations, which occur at Ser31, Ser32, or Thr35 (**Figure S6**). mTlaA2 was also digested with endoproteinase LysC, resulting in fragments **3**, **4**, and **5** (**Figure 3D**). Fragment **3** had been dehydrated four times and contained one ring and three dehydro amino acids. Fragment **4** had been dehydrated once, and product **5** twice (**Figure 3D**), suggesting that of the three Ser and Thr residues in peptide **5** (Ser31, Ser32, and Thr35; numbering based on core peptide, **Figure 2**), two are dehydrated.

Collectively, the MS data on TlaM-modified TlaA1 and TlaA2 show that they generally produce very similar products. Whereas TlaA2 is cleanly dehydrated seven times, TlaA1 differs in that Ser3 is not dehydrated in the six-fold dehydrated product and Ser3 and Thr4 are not dehydrated in the five-fold dehydrated product. The lack of dehydration of both Ser3 and Thr4 in the five-fold dehydrated product was shown by endoproteinase AspN digestion that demonstrates that all five dehydrations are in the segment Asp6-Cys36 (see bioactivity section). Several possible explanations can be offered for the difference between TlaA1 and TlaA2. First, the Thr8 and Thr9 residues in TlaA2 corresponding to Ser3/Thr4 in TlaA1 are further removed from the end of the leader peptide. It has been demonstrated previously for other class II synthetases that a minimal distance is required between the leader peptide residues that are involved in binding the enzyme and the first residue that can be dehydrated efficiently.[^59^] Thus, Thr8/Thr9 may be dehydrated more efficiently than Ser3/Thr4 because they can reach the active site better. We suspect that Ser3/Thr4 may not be dehydrated at all in the native setting because Ser3/Thr4 are in the heptapeptide that we hypothesize is removed in a second proteolytic step and for which we provide support in the bioactivity section. The reason that these residues are dehydrate at all in *E. coli* is likely because TlaM is expressed from a high copy plasmid (pRSFDuet; ∼100 copies) and TlaA1 is expressed from a lower copy plasmid (pETDuet; ∼40 copies). Hence, the concentration of enzyme is likely much higher in this synthetic biology study than in the native organism.

Based on the tandem MS data, both peptides are likely to contain identical ring patterns (Figure 3E) with two non-overlapping rings near the N-terminus (ring A) and in the center of the peptide (ring B) and two overlapping rings C and D at the C-termini. The tandem MS data did not allow determination which of the three Ser/Thr residues near the C-terminus were involved in ring formation, nor did the MS data conclusively determine whether ring A contains five amino acids (as shown in Figure 3E) or four amino acids.

### 2.4 Determination of ring pattern by NMR spectroscopy

Collectively, the proteolytic digests and tandem MS analysis established the presence of a MeLan ring between former Thr18 and Cys23 in mTlaA1 and between former Thr22 and Cys27 in mTlaA2 (rings B). Since no fragmentation was observed in the C-terminal region of both mTlaA1 and mTlaA2, overlapping ring patterns are likely formed from the sequence SSLATPCKRC in which two of the three Ser/Thr residues were dehydrated (**Figure S5** and **Figure S6**). Because tandem MS analysis cannot deduce the ring pattern of these overlapping rings,[^58^] fragment **5** was subjected to NMR analysis (**Figure 4A**). The ring pattern of the 10-amino acid peptide **5** that was cleanly dehydrated twice was elucidated using 1D and 2D NMR experiments, including ^1^H, ^1^H-^1^H TOCSY, ^1^H-^1^H NOESY, and ^1^H-^13^C HSQC. The ^1^H and ^13^C chemical shift assignments of the peptide in 90% H_2_O and 10% D_2_O at 25 °C are summarized in **Table S3**. A total of 7 amide protons were observed (**Figure S7**). The missing NH protons correspond to the N-terminal residue Ser1 (residue numbering based on the LysC product **5**, **Figure 4A**), the residue at position 2 (formerly Ser2), and a proline at position 6. Both the ^1^H and ^13^C chemical shift values of the former Ser2 residue and former Thr5 showed significant upfield shifts compared to typical Ser and Thr residues, suggesting their involvement in lanthionine and methyllanthionine formation (**Figure S8**). These changes include β-proton signals at 3.21 ppm and 3.04 ppm, and a β-carbon at 33.8 ppm for the former Ser2. The β-proton and β-carbon of Thr5 shifted to 3.64 ppm and 40.8 ppm, respectively, compared to the typical values of approximately 4.10 ppm (^1^H) and 69.0 ppm (^13^C) observed when bonded to a more electronegative oxygen atom. Further analysis of the NOESY data suggested that a lanthionine was formed between Ala2 and Cys7, and a methyllanthionine was formed between Thr5 and Cys10. This assignment is supported by a set of NOE cross-peaks observed between these residues across the thioether bond. For example, a strong NOE was detected between the β-protons of former Cys7 and Ala2 (former Ser2) (**Figure 4B**). Similarly, NOEs were observed between the amide proton of former Cys10 and the β-proton of former Thr5, and between the amide proton of former Thr5 and the β-proton of former Cys10 (**Figure 4C**).

**Figure 4.**
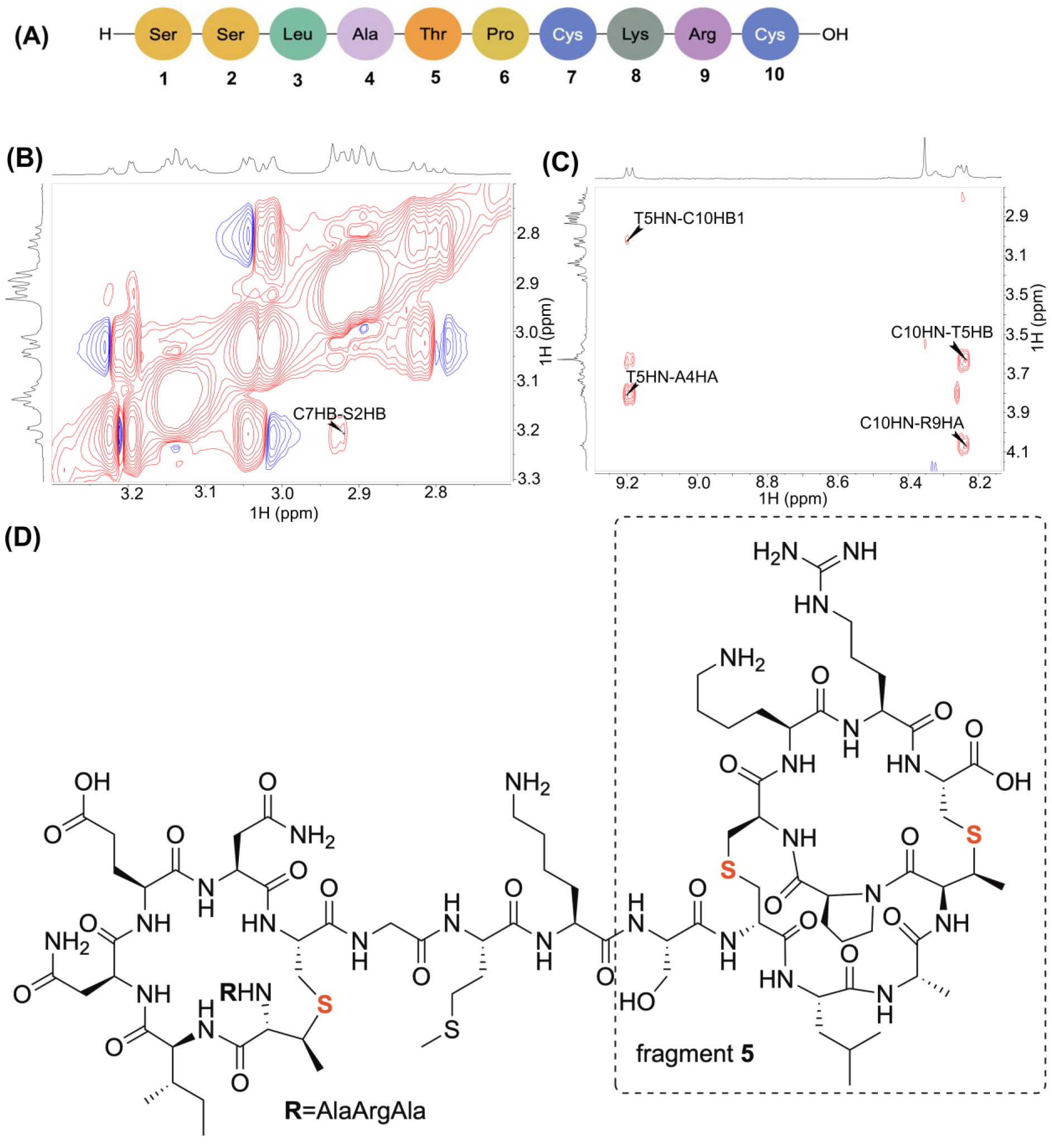
NMR analysis of fragment 5. A) Sequence of fragment **5** and residue numbering used in the description of the NMR data. B) ^1^H-^1^H NOESY spectrum of fragment **5**. Cross-peaks across the lanthionine bond are observed between the β-protons of former Ser2 and former Cys7. C) ^1^H-^1^H NOESY spectrum of fragment **5**. Cross-peaks across the methyllanthionine bond are observed between the amide proton of former Thr5 and the β-proton of former Cys10, and vice versa between former Cys10 and Thr5. D) Fragment **2** showing the ring patterns of rings B, C, and D. Fragment **5** is boxed.

As noted above, the amide proton of Ala2 (former Ser2) was absent in the TOCSY spectrum, and no NOE cross-peaks were observed between Leu3 and Ala2, likely due to line broadening. These missing peaks prevented an unambiguous distinction between ring formation between former Cys7 and either the first or second residue in fragment **5**. To resolve this ambiguity, the longer 22-amino acid peptide, fragment **2** derived from mTlaA1, was used for further analysis (**Table S4**). In this peptide, the two former serine residues in question are positioned at positions 13 and 14 and were clearly assigned (**Figure S9**). A distinct NOE cross-peak was observed between the amide protons of residue 14 and Leu15 (**Figure S10**, corresponding to Ala2 and Leu3 in fragment **5**), as well as between the amide proton of residue 14 and the β-proton of Ser13 (**Figure S11**, corresponding to Ala2 and Ser1 in fragment **5**). Therefore, the former Ser2 was assigned as the dehydrated serine in fragment **5**. Collectively, the proteolytic cleavage fragments combined with NMR analysis confirmed the ring patterns in mTlaA1 and mTlaA2 for rings B, C, and D (**Figure 4D**).

### 2.5 Stereochemistry of the (methyl)lanthionines in mTlaA1 and mTlaA2

The stereochemical configurations of the rings in mTlaA1 and mTlaA2 were determined by Marfey analysis.[^60^] mTlaA1 (mixture of 5 and 6-fold dehydrated peptide) and mTlaA2 (7-fold dehydrated) were used since the tandem MS data shows that the heterogeneity is caused by partial dehydration of Ser3 and Thr4, which are not involved in ring formation. The peptides were hydrolyzed in acid followed by reaction with the advanced Marfey’s reagent, Nα-(5-fluoro-2,4- dinitrophenyl)-L-leucinamide (L-FDLA).[^61^] The resulting derivatized products were compared with authentic standards using LC-MS. The standard used for lanthionine analysis was CylL_L_, which contains DL-Lan, LL-Lan, and LL-MeLan.[^29^] Based on the retention times and co-injection with the standards, mTlaA1 contains DL-lanthionine (**Figure 5A**). The standards for methyllanthionine were obtained from mCoiA1, which contains D-*allo*-L-MeLan, LL-MeLan, and DL-MeLan.[^62^] The L-FDLA-derivatized MeLan from hydrolyzed mTlaA1 coeluted with DL-MeLan, which confirmed that the methyllanthionine rings all have DL-stereochemistry (**Figure 5B**). This observation was unexpected because the N-terminal sequence has a common motif (Dhx-Dhx-Xxx-Xxx-Cys; Dhx = Dha or Dhb) that resulted in LL-(Me)Lan in other lanthipeptides.[^29, 63–65^] Similar analysis of mTlaA2 indicated that it also contained DL-Lan and DL-MeLan (**Figure 5C,D**).

**Figure 5.**
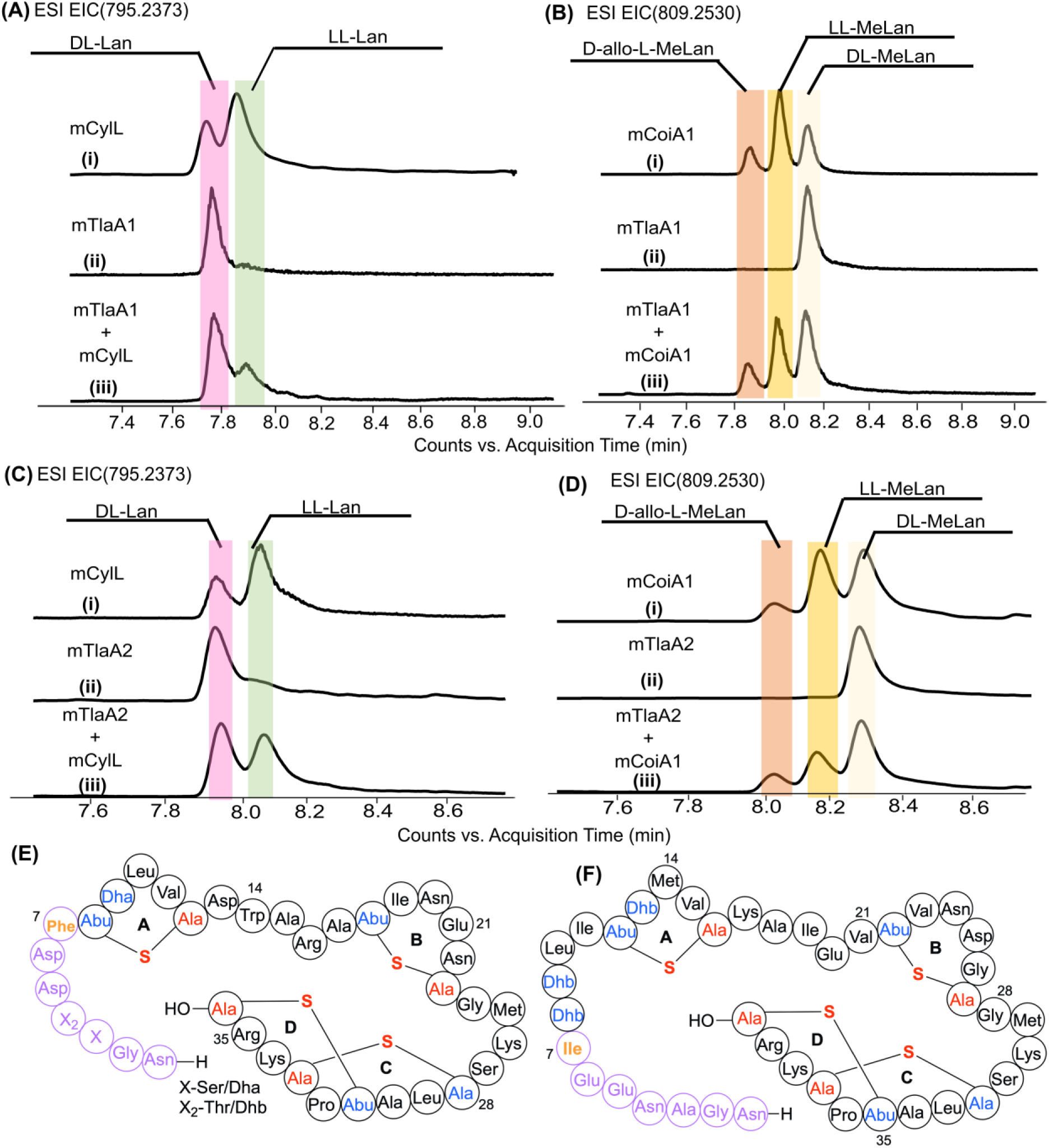
Hydrolyzed peptides were derivatized with Marfey’s reagent and analyzed by LC–MS to determine the stereochemistry of thioether crosslinks. Extracted ion chromatograms (EICs) of L-FDLA derivatized mTlaA1, mTlaA2, MeLan standard (from mCoiA1), and Lan standard (from CylL_L_). **A, B.** EICs corresponding to m/z 795.2373 and 809.2530 are shown for Marfey-derivatized lanthionine (Lan) and methyllanthionine (MeLan), respectively, of modified TlaA1. **C, D.** EICs for modified TlaA2. (i) MeLan or Lan standard, (ii) tested compound, and (iii) coinjection (tested compound + standard). **E.** The proposed structure of mTlaA1 after LahT150 treatment; Abu, 2-aminobutyric acid. **F.** The proposed structure of mTlaA2 after LahT150 treatment. Highlighted amino acid residues in red and blue are where PTMs occurred. The residues in pink and orange would be removed if a protease cleaves off the N-terminal 6 or 7 residues, as discussed in the text.

These data did not distinguish between a Lan or a MeLan as the A ring in mTlaA1 (involving Cys12 and either Thr8 or Ser9), nor whether the A-ring in mTlaA2 is formed between Cys16 and Thr12 or Thr13. To determine the residues involved in forming the N-terminal ring A, we also subjected fragment **1** derived from mTlaA1 to acidic hydrolysis and Marfey’s analysis. We confirmed that the MeLan A-ring in mTlaA1 has a DL configuration, and not an LL-configuration (**Figure S12**). But surprisingly, we also observed DL-Lan, albeit with a signal intensity that is about 5-fold smaller than that observed for MeLan. These findings suggest that in mTlaA1, both a MeLan is formed between Cys12 and Thr8 and a Lan is formed from Cys12 and Ser9. Lanthipeptide synthases usually do not form alternative ring patterns, and when other patterns are observed, they usually arise from non-enzymatic ring formation that competes with enzymatic ring formation. Because non-enzymatic MeLan formation is much slower than non-enzymatic Lan formation,[^12, 66^] and because the stereochemistry of MeLan formation from the Dhb-Dha-Leu-Val-Cys motif is opposite to that expected from non-enzymatic cyclization (and therefore is likely to require enforcement by the enzyme),[^67^] we hypothesize that the small amount of Lan formed from Cys12 and Ser9 is non-enzymatic and non-physiological. Collectively, the data in this study suggest the structures shown in **Figure 5E** and **F** for the lanthipeptides from *Thermoactinomyces sp.* DSM 45891.

### 2.6 Antimicrobial activity

We next evaluated the antimicrobial activity of mTlaA1 and mTlaA2 cleaved with LahT150 and dissolved in 5% DMSO. The proteolytic product derived from mTlaA1 showed very weak antimicrobial activity against *Bacillus subtilis* and *E. coli* (**Figure S13**), whereas the product from mTlaA2 showed no detectable antimicrobial activity. When the two modified lanthipeptides were combined to assess potential synergistic activity, no antimicrobial activity was observed. In other two-component lantibiotics such as haloduracin,[^68^] cytolysin,[^69, 70^] and lichenicidin,[^22^] a second proteolytic step is involved after removal of the leader peptide by the PCAT. In all three cases, the N-terminal sequence that is removed is a hexapeptide. For geobacillin II, a heptapeptide has been suggested to be removed in such a second proteolytic step.[^47^] In some cases, the required second protease is encoded within the BGC, but in many other cases, the second protease is encoded elsewhere.[^14, 71^] Because the *tla* BGC does not encode a second protease, we used AspN endopeptidase cleavage of mTlaA1 to access a 31-mer, **6,** with 5 dehydrations and including rings A-D (**Figure 6A**). Peptide **6** demonstrated antibacterial activity against *B. subtilis* and the Gram-negative ESKAPE pathogens *Acinetobacter baumannii* and *Klebsiella pneumoniae* (**Figure 6B-D**). Growth inhibition of Gram-negative bacteria by lanthipeptides is a rare observation.[^72^] These findings suggest that a second proteolytic step is likely required for bioactivity, although the precise site is currently not known and peptide **6** is unlikely to be the native peptide. The two-step proteolytic activation process unfortunately resulted in quantities too small to determine minimal inhibitory concentrations, with especially the second proteolysis step with AspN proving low-yielding, possibly because the cleavage side is near a D-stereocenter associated with the DL-MeLan A-ring. In agreement with the hypothesis presented in section 2.3, the observed bioactivity with peptide **6** shows that the two inefficient dehydrations in mTlaA1 at Ser3 and Thr4 are not required for bioactivity. We termed the active peptide **6,** thermolanthin A, and anticipate that the natural peptide will likely have higher activity. We tried to perform a similar sequence of two proteolytic events on mTlaA2. Unfortunately, treatment with endoproteinase GluC not only resulted in cleavage at Glu6 but also at Glu20 in the core peptide.

**Figure 6.**
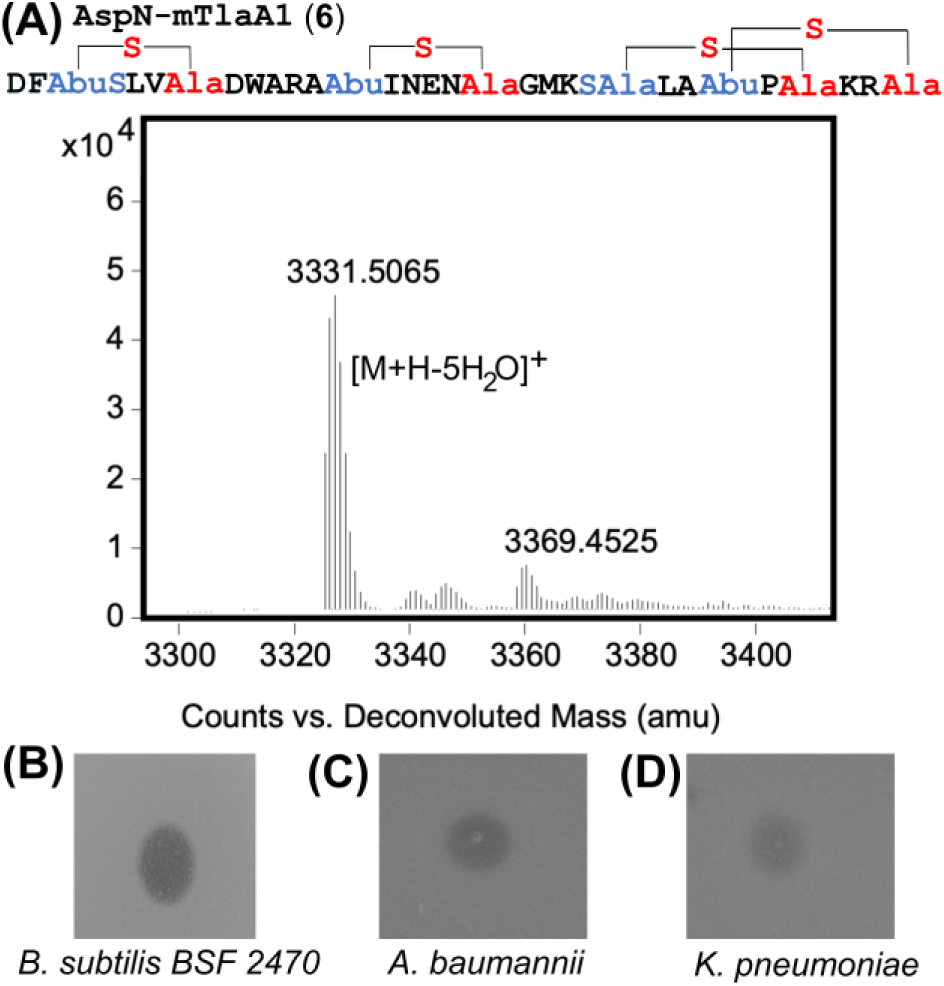
ESI-MS characterization and bioactivity assessment of AspN-digested mTlaA1. A) ESI-MS spectrum showing the molecular mass of AspN-digested mTlaA1. Observed m/z [M+H-5H_2_O]^+^ 3331.5065, calculated 3330.5097. B) Bioactivity assay against *B. subtilis* 2470 and two Gram-negative ESKAPE pathogens. For each assay, 2 µL of AspN-cleaved mTlaA1 (∼500 µM) was spotted.

## 3. Conclusion

This work identified two new lanthipeptides from the thermophile *Thermoactinomyces sp.* DSM 45891. The thioether patterns of these peptides shown in **Figure 5E** and **5F** have not been previously reported for known lanthipeptides.

The two peptides are very similar in structure and AspN-processed mTlaA1 displayed antibacterial activity without the need for a partner peptide. We therefore do not have any evidence that the *tla* BGC generates a two-component lanthipeptide. Based on the current data, it is more likely that the two peptides are congeners. In the absence of information of the products formed by the producing organism, the precise proteolytic processing site remains unresolved. It is likely that the products of the *bai* and *cro* BGCs will have a similar ring pattern to thermolanthin A with respect to rings B-D, which could be the bioactivity-inducing part of these molecules.

This study showed that TlaM catalyzes the stereoselective formation of both DL-MeLan and DL-Lan rings during post-translational modification of its precursor peptides. Based on previously observed stereochemical selectivity with Dhx-Dhx-Xxx-Xxx-Cys motifs in which LL-Lan and MeLan structures are usually generated as a result of an inherent preference imparted by the motif,[^29, 63, 64, 67^] the formation of a DL-MeLan A-ring is unique and suggests TlaM enforces the formation of a DL-MeLan. The mechanism by which it achieves this selectivity requires further investigation.

## 4. Experimental Section

### 4.1 General Experimental Procedures and Materials

Molecular cloning was performed using the primers and gblock listed in Supporting Information **Table S5** and **Table S6**, respectively, to produce three different plasmids: His6-SUMO-TlaA1-pETDuet, His6-SUMO-TlaA2-pETDuet, His6-SUMO-TlaA2E20D-pETDuet, His6-SUMO-TlaA2E6D-pETDuet, and His6-TlaM-pRSFduet. Co-expression of His_6_-SUMO-TlaA1 and His_6_-TlaM-pRSFduet was carried out in *E. coli* BL21(DE3). Similarly, co-expression of His6-SUMO-TlaA2 and His6-TlaM-pRSFduet was done in the same expression strain. Purification by NiNTA affinity chromatography was performed after expression, followed by proteolytic cleavage with LahT150, and MALDI-TOF-MS analysis using a Bruker Ultraflex II. Cleaved peptide was desalted using a C18 ziptip and eluted in a saturated solution of super-DHB in 80% MeCN, which served as MALDI matrix. The cleaved peptide was stored at −20 °C pending further purification.

### 4.2 Co-expression using His6-SUMO-TlaA-pETDuet and His6-TlaM-pRSFduet

For each TlaA/TlaM co-expression, 4 L flasks containing 1 L of LB medium with 50 µg/mL kanamycin and 100 µg/mL ampicillin were inoculated with 10 mL of an LB overnight culture of *E. coli* BL21(DE3) carrying the respective expression plasmids. Cells were grown at 37 °C until the cultures reached an optical density at 600 nm (OD_600_) of ∼0.75-0.95. Then, the culture flasks were placed on ice for 30 min, and the shaker temperature was lowered to 18 °C. Next, 0.5 mL of sterile stock solution of isopropyl β-D-1-thiogalactopyranoside (IPTG, 1 M) was added to each flask. After overnight incubation at 18 °C, cultures were harvested by centrifugation (6000×g, 4 °C, 10 min).

The cell pellets were resuspended in lysis buffer (20 mM Na_2_HPO_4_, 10 mM imidazole, 300 mM NaCl and 6 M guanidine hydrochloride, pH 7.5) and lysed using a sonicator (on 4 s and off 3 s) at 50 % amplitude. Afterwards, the lysate was cleared by centrifugation (10 000×g, 4 °C, 30 min) and filtered using a syringe filter (0.45 µm). Peptide isolation was performed by NiNTA affinity chromatography using pre-equilibrated 0.5 mL/ L of expression of NiNTA resin in a gravity-flow column. After applying the cell lysate, the column was washed with 3 mL of lysis buffer, followed by 3 mL of wash buffer 1 (20 mM Na_2_HPO_4_, 10 mM imidazole, 300 mM NaCl and 4 M guanidine hydrochloride, pH 7.5) and wash buffer 2 twice (20 mM Na_2_HPO_4_, 50 mM imidazole, 300 mM NaCl, pH 7.5). Next, the peptide was eluted from the resin by adding 3 mL of elution buffer (20 mM Na_2_HPO_4_, 1 M imidazole, 300 mM NaCl, pH 7.5).

### 4.3 Purification of His6-LahT150

Similar protein expressions using His6-LahT150-pET15 were carried out as described for the peptide co-expressions above. The cells were harvested by centrifugation (5 000×g, 4 °C, 15 min), and cell pellets were resuspended in protein Start buffer (20 mM Tris-HCl, 10 mM imidazole, 300 mM NaCl and 10% glycerol, pH 8.0) containing 4 mg/mL of lysozyme. This mixture was incubated at 4 °C, then further lysed via sonication (40% amplitude, 4.4 s on, 5.5 off) for 6 min. The lysate was centrifuged at 10,000 x g for 30 min. The supernatant was loaded onto an immobilized metal affinity chromatography (IMAC) column equilibrated with protein Start buffer, using 0.5 mL of Ni resin per gram of cell pellet. This step was followed by adding 2–3 column volumes (CV) of protein Start buffer to the column. The column was then washed with 2-3 CV of protein Wash buffer (20 mM Tris-HCl, 30 mM imidazole, 300 mM NaCl and 10% glycerol, pH 8.0) before eluting the protein with 2-3 CV of protein Elution buffer (20 mM Tris-HCl, 300 mM imidazole, 300 mM NaCl and 10% glycerol, pH 8.0). Each fraction from the wash and elution was collected for analysis by sodium dodecyl sulfate polyacrylamide gel electrophoresis (SDS-PAGE) using a 4–20% Tris gel at 150 V. The elution fraction containing LahT150 was collected, and the buffer was exchanged to protein storage buffer (20 mM Tris-HCl, 500 mM NaCl, and 10% glycerol, pH 8.0) with a 3 kDa MWCO Amicon filter. Subsequently, the fraction was aliquoted and stored at −80 °C.

### 4.4 Protease Digestion

Proteolytic cleavages were performed using the proteases LahT150, chymotrypsin, LysC, GluC, and AspN endopeptidase under standard digestion conditions. Purified peptides (1-3 mg/mL) were dissolved in the appropriate buffer compatible with each protease (pH 7.8–8.0). Proteases were added at an enzyme-to-substrate ratio of 1:100. Digestions were carried out at 37 °C for all proteases except LahT150, which was carried out at RT for 16-18 h. Reactions were quenched by acidification with formic acid to a final concentration of 0.1–1% (v/v). The digested samples were analyzed directly by MALDI-TOF-MS to verify proteolytic cleavage. Next, the digested products were further purified by HPLC for further analysis.

### 4.5 HPLC purification of TlaA-derived product peptides

The protease-digested peptides were purified by preparative HPLC using an Agilent 1260 HPLC system. For separation, a Luna 10 µm C5 100 Å, LC Column 250 x 21.2 mm was used, employing a solvent system consisting of H_2_O/0.1% trifluoroacetic acid (solvent A) and MeCN/0.1% trifluoroacetic acid (solvent B) and a linear gradient of 2-100% B in 45 min. Peptide fractions were identified by MALDI-TOF-MS, pooled, and the solvent was removed using a lyophilizer.

### 4.6 NEM Assay of LahT150-digested mTlaA1 and mTlaA2

A 25-μL reaction was prepared with 25-50 μM peptide dissolved in 50 mM Tris-HCl, pH 7.5 (12.5 μL), along with 0.3 mM of fresh 1.0 mM TCEP stock solution (10 μL). The mixture was incubated at room temperature for 30 min before adding 2.5 μL of 500 mM NEM. The reaction was then incubated at room temperature for 3 h, desalted using a C18 ZipTip, and analyzed by MALDI-TOF MS. The formation of an adduct is indicated by a mass shift of +125 Da.

### 4.7 DTT Assay of LahT150-digested mTlaA1 and mTlaA2

Modified TlaA1 and TlaA2 that had been cleaved by LahT150 were dissolved in 50 μL of 100 mM Tris-HCl buffer, pH 7.5, to obtain a final concentration of 100 μM. *N*, *N*-diisopropylethylamine (DIPEA) and DTT were added to give final concentrations of 400 mM and 500 mM, respectively. The 50 μL reaction was incubated at room temperature for 24 h. After desalting with a C18 ziptip, the reaction was analyzed by MALDI-TOF MS.

### 4.8 Tandem Mass Spectrometry

The HPLC-purified protease cleaved precursor peptides were injected onto the Agilent 6545B Quadrupole Time-of-Flight mass spectrometer. Samples were run on a Phenomenex Cat. No. 00F-4497-AN Kinetex C800 column (2.6 µm, 100 Å, 150 x 2.1 mm) using a solvent system consisting of H_2_O/0.1% formic acid (solvent A) and MeCN/0.1% formic acid (solvent B) and a linear gradient of 5-100% B over 20 min at a flow rate of 0.4 mL/min. The parameters used for MS(/MS) data acquisition included a resolution of 30,000 in positive mode, an isolation width of 2-3 Da, and a normalized collision energy of 20, 25, and 30 (MS/MS). Data analysis was performed using MassHunter Qualitative Analysis.

### 4.9 Marfey’s Analysis of mTlaA1 and mTlaA2

For the Marfey analysis,[^60^] mCylL_L_ was used for the lanthionine standard and mCoiA1 for the methyllanthionine standard. The standards, along with modified TlaAs, were purified after heterologous expression in *E. coli* as described previously, followed by acidic hydrolysis, and the Lan and MeLan were derivatized. Dried peptides (100 μg) were dissolved in 0.8 mL of 6 M DCl in D_2_O for hydrolysis using a long screw glass vial. N_2_ gas was bubbled into the glass vials for 2 min. The reaction was heated to 120 °C and stirred for 20 h. Each reaction product was then dried using a rotary evaporator and placed in the lyophilizer for 45 min to remove any remaining DCl. After drying the samples, 600 μL of 0.8 M NaHCO_3_ (H_2_O) and 400 μL of 10 mg/mL of Nα-(5-fluoro-2,4-dinitrophenyl)-l-leucinamide (L-FDLA) in acetonitrile were added to the glass vials. The mixture was vortexed, and the glass vials were stirred in the dark at 67 °C for 3 h. Next, 100 μL of 6 M HCl was added to the derivatization product, and the mixture was vortexed. The mixture was placed in the lyophilizer to dry, then resuspended in 400 μL of acetonitrile and transferred to a 1.5 mL centrifuge tube. The resuspension mixture was centrifuged for 10 min, and the supernatant was collected and analyzed on an Agilent 6545 LC/Q-TOF instrument using a Kinetex 1.7 μm F5 100 Å, LC column (100 × 2.1 mm; Phenomenex; part no.00D-4722-AN) at a constant flow rate of 0.4 mL/min, and a gradient of 5-80 % of Solvent B (MeCN with 0.1% formic acid) in Solvent A (water with 0.1% formic acid). The following exact masses were extracted for MeLan (m/z 809.25299) and Lan (m/z 795.23734) from the chromatographic separation of each derivatized product and the co-injections.

### 4.10 NMR analysis

Fragments **2** and **5** were dissolved in 90% H_2_O and 10% D_2_O. Based on peak intensities, the estimated concentrations were ∼0.23 mM for fragment **5** and ∼0.20 mM for fragment **2**. All NMR data were collected at 25 °C on a Bruker Avance NEO 600 MHz spectrometer equipped with a 5-mm curio BBO prodigy probe, including one-dimensional (1D) and two-dimensional (2D) homonuclear ^1^H-^1^H TOCSY (total correlation spectroscopy which reveals the correlation of protons in the same spin system) at 80 ms mixing time, 2D ^1^H-^1^H NOESY (Nuclear Overhauser Effect spectroscopy which reveals close spatial proximity between two protons) at 400 ms mixing time, as well as a 2D multiplicity edited ^1^H-^13^C HSQC (heteronuclear single quantum coherence spectroscopy, revealing one-bond correlation between ^1^H and ^13^C in which CH/CH_3_ cross-peaks appear with opposite phase relative to CH_2_ groups). All spectra were acquired using standard pulse programs in Bruker Topspin 4.1.4 software and processed and analyzed using Mnova (version 15.1.0.; Mestrelab Research).

### 4.11 Agar Well Diffusion Assays to Determine Antimicrobial Activity

Bioassays were performed using agar well-diffusion growth-inhibition assays with the protease-cleaved purified peptides. Peptides were dissolved in 50 mM Tris-HCl, 100 mM NaCl, pH 7.5 (initial assay peptide dissolved in 5 % DMSO) to achieve a starting concentration of ∼0.5-1.8 mM. Agar plates were prepared by making a bottom layer with LB growth medium supplemented with 1.5% (w/v) agar (20–25 mL per 90 mm plate). The agar was allowed to solidify at room temperature. Next, the soft agar was prepared by mixing an overnight culture diluted to ∼10⁶–10⁷ CFU/mL with molten LB agar containing 0.7% (w/v) agar. The mixture was poured over the solidified base layer and allowed to solidify. Then 1.5 μL of the positive controls, kanamycin (4.29 mM) or bacitracin (35 mM), was added to each plate as well as 2 μL of the tested compounds. The dried plates were incubated at 37 °C for 18 h, and the presence or absence of zones of growth inhibition was used to determine activity. The negative control was 50 mM Tris-HCl, 100 mM NaCl, pH 7.5.

## Funding

This manuscript is the result of funding in whole or in part by the National Institutes of Health (grant R01 AI144967 to W.A.V.) and therefore it is subject to the NIH Public Access Policy. Through acceptance of this federal funding, NIH has been given a right to make this manuscript publicly available in PubMed Central upon the Official Date of Publication, as defined by NIH. A Bruker UltrafleXtreme mass spectrometer used was purchased with support from the Roy J. Carver Charitable Trust (Grant No. 22-5622). W.A.V. is an Investigator of the Howard Hughes Medical Institute.

## Conflicts of interest

The authors declare no conflicts of interest.

## Data availability Statement

All data in this study are available at: Weir, Enleyona; Zhu, L; van der Donk, Wilfred (2026), “Data associated with “Structure and activity of a class II lanthipeptide from a thermophilic bacterium””, Mendeley Data, V1, doi: 10.17632/jgbnmfyx44.1

## Additional Information

The following files are available free of charge:

Supporting Information with additional figures, tables with primers and chemical shift assignments.

## Notes

### Competing Interest Statement

The authors have declared no competing interest.

### Summary of Updates

New text was added to place the current work in context of other lanthipeptide BGCs that have multiple precursors. Text was added to offer explanations for why TlaM processes the two peptides differently. Panel E was added to Fig 3,

