## Supporting Information for "Structure and Activity of Class II Lanthipeptides from a Thermophilic Bacterium"

Table S1: Accession number of proteins used ........................................................S2
Table S2: Calculated and observed masses for mTlaAs in Figure 3 ........................S2
Table S3. $^{1}$H and $^{13}$C chemical shift assignments of fragment **5** ...............................S3
Table S4. $^{1}$H and $^{13}$C chemical shift assignments of fragment **2** ............................S3-5
Table S5: Primers used in this study .........................................................................S5
Table S6: Codon-optimized genes used in this study ............................................S5-6
Figure S1. Sequence alignment of TlaM and homologs ............................................S7
Figure S2. EIC of dehydration patterns of mTlaAs .....................................................S8
Figure S3. GluC/LysC cleavage of mTlaA1..................................................................S9
Figure S4. Tandem MS of fragment **A** .......................................................................S10
Figure S5. Tandem MS of fragment **B** .......................................................................S11
Figure S6. Tandem MS of AspN cleaved mTlaA2......................................................S12
Figure S7. $^{1}$H-$^{1}$H TOCSY spectrum of fragment **5**....................................................S13
Figure S8. $^{1}$H-$^{13}$C HSQC spectrum of fragment **5** ....................................................S14
Figure S9. $^{1}$H-$^{1}$H TOCSY spectrum of the fragment **2**.............................................S15
Figure S10: The amide region of the $^{1}$H-$^{1}$H NOESY spectrum of fragment **2**.........S16
Figure S11. The amide region of the $^{1}$H-$^{1}$H NOESY spectrum of fragment **2**.........S17
Figure S12. Marfey's analysis of fragment **1** ................................................... S18-S19
Figure S13. Bioactivity assay of mTlaAs ...................................................................S20
References..................................................................................................................S21

**Table S1**: **Accession number of protein used**

| | |
|---|---|
| WP_072335371.1 | TlaA1 |
| WP_072335368.1 | TlaA2 |
| WP_177239804.1 | TlaM |
| WP_072335376.1 | TlaT |
| WP_131847683.1 | BaiD |
| WP_131847685.1 | BaiA |
| WP_131847687.1 | BiaM |
| WP_131847689.1 | BaiT |
| WP_131847692.1 | BaiH |
| WP_307253290.1 | CroH |
| WP_307253292.1 | CroT |
| WP_307253293.1 | CroM |
| WP_307253294.1 | CroA |
| WP_307253295.1 | CroD |

**Table S2.** Calculated and observed masses for mTlaAs in Figure 3.

| | |
|---|---|
| Fig. 3A | mTlaA1 observed $[M+H-5\ H_2O]^+$ m/z=3805.7; calculated=3804.7; $[M+H\text{-}6H_2O]^+$ m/z=3787.7; calculated=3786.7; $[M+H-7\ H_2O]^+$ m/z=3770.9; calculated m/z=3668.7. DTT adduct observed, $[M+H-5\ H_2O+1\ DTT]^+$ m/z=3959.5; calculated m/z=3958.7; $[M+H-6\ H_2O+1\ DTT]^+$ m/z=3941.5; calculated m/z=3940.7. A mass change of +308 Da for two DTT adducts, observed $[M+H-6\ H_2O+2\ DTT]^+$ m/z=4095.5; calculated m/z=4095.7. |
| Fig. 3B | ESI-MS (positive mode) of fragment **1** showed a prominent ion at $[M+H\text{-}2\ H_2O]^{3+}$ m/z=1227.2370, calculated m/z=1226.9897; $[M+H-3\ H_2O]^{3+}$ m/z=1220.9037, calculated m/z=1220.9846. Fragment **2** showed $[M+H-3\ H_2O]^{4+}$ m/z=576.0306, calculated m/z=576.1945. |
| Fig. 3C | mTlaA2, observed $[M+H-7\ H_2O]^+$ m/z=4077.0; calculated m/z=4075.9. Mass changes of +308 Da and +462 Da were observed, respectively, $[M+H-7\ H_2O+2\ DTT]^+$ m/z 4385.3; calculated 4383.9; $[M+H-7\ H_2O+3\ DTT]^+$ m/z=4538.2; calculated m/z=4537.9. |
| Fig. 3D | ESI-MS (positive mode) of fragment **3** showed a $[M+H-4\ H_2O]^+$ m/z=1765.8692, calculated m/z=1765.8612; **4** $[M+H-\ H_2O]^+$ m/z=1318.6101, calculated m/z=1318.6130. Fragment **5** showed $[M+H-2\ H_2O]^+$ m/z =1029.4960, calculated m/z=1029.4968. |

**Table S3**. $^{1}H$ and $^{13}C$ chemical shift assignments of fragment **5** in 90% $H_2O$ and 10% $D_2O$ at 25 °C

| # | AA ID | N-H | αH | βH | γH | δH | other |
|---|---|---|---|---|---|---|---|
| 1 | S | - | 3.62<br>55.5 | 3.74, 3.70<br>63.5 | | | |
| 2 | (S) A | NA | 4.51<br>53.6 | 3.21, 3.04<br>33.8 | | | |
| 3 | L | 7.90 | 4.33<br>51.6 | 1.58<br>40.0 | 1.48<br>23.9 | 0.83,<br>22.2<br>0.77,19.8 | |
| 4 | A | 8.26 | 3.81<br>51.7 | 1.30<br>14.9 | | | |
| 5 | (T)<br>Abu | 9.19 | 4.56<br>56.3 | 3.64<br>40.8 | 1.1718.9 | | |
| 6 | P<br>(trans) | - | 4.33<br>61.8 | 2.26, 1.99<br>29.8 | 2.10,<br>1.98<br>24.6 | 3.62,<br>3.55<br>48.0 | |
| 7 | C | 7.71 | 4.54<br>53.7 | 2.93<br>34.7 | | | |
| 8 | K | 7.15 | 4.28<br>52.7 | 1.78, 1.64<br>31.3 | 1.28<br>21.8 | 1.58<br>26.8 | 2.91<br>39.6 |
| 9 | R | 8.32<br>(br) | 4.07<br>55.1 | 1.73<br>27.5 | 1.65<br>24.3 | 3.14<br>40.8 | |
| 10 | C | 8.24 | 4.40<br>58.7 | 3.03, 2.81<br>32.3 | | | |

NA: not observed

NOE observed between NH of Thr5 and Hβ of Cys10, and between NH of Cys10 and Hβ of Thr5, between Hβ protons of Thr5 and Cys10.

NOE observed between Hβ of Cys7 and Hβ of Ala2, between Hβ protons of Cys7 and Ala2.

Serines converted to Lan are shown as (S) A; Thr residues converted to MeLan are shown as (T) Abu. Pairs of residues that are crosslinked are shown in the same color.

**Table S4**. $^{1}H$ and $^{13}C$ chemical shift assignments of fragment **2** in 90% $H_2O$ and 10% $D_2O$ at 25 °C

ARATINENCGMKSSLATPCKRC (peptide in red = peptide **5**)

| # | AA ID | N-H | αH | βH | H | ph. | other |
|---|---|---|---|---|---|---|---|
| 1 | A | | | | | | |
| 2 | R | | 4.165 | 1.53 | 1.18, 1.12 | 1.53 | 2.82 |
| 3 | A | | | | | | |
| 4 | (T) Abu | 8.129 8.168 8.368 | 4.61 | 3.57 (CH) 43.5 | 1.235 | | |
| 5 | I | 7.727 | 4.24 | 1.92 | 1.22 1.05 | 0.78 | |
| 6 | N | 8.343 7.93 | 4.50 | 3.11, 2.93 | | | |
| 7 | E | | | | | | |
| 8 | N | | 4.55 | 2.83 | | | |
| 9 | C | 7.88 | 4.357 | 2.95 | | | |
| 10 | G | | | | | | |
| 11 | M | 8.35 | | | 2.82 | 2.02 (CH3) | |
| 12 | K | | 4.165 | 1.53 | 1.18, 1.12 | 1.53 | 2.82 |
| 13 | S | 8.126 | 4.169 | 3.85, 3.75 61.2 | | | |
| 14 | (S) A | 7.35 | 4.48 | 3.21, 3.09 | | | NOE from NH (7.45) to Ha (4.169) of S13. NOE from NH (7.35) to L15 (7.66) and to S13 (8.126) |
| 15 | L | 7.66 | 4.30 | 1.55, 1.44 39.8 | 1.55 26.5 | 0.81 22.0 | |
| 16 | A | 8.314 | 3.79 51.6 | 1.288 | | | |
| 17 | (T) Abu | 9.184 | 4.561 | 3.644 4.08 | 1.165 | | involve -S- NOE: NH- 3.03 |
| 18 | P (trans) | | 4.30 | 2.26, 1.99 29.8 | 2.06 | 3.57 48.2 | |

| | | | | | | | |
|---|---|---|---|---|---|---|---|
| 19 | C | 7.76 | 4.49 | 2.89 | | | NOE: HB-3.21 |
| 20 | K | 7.09 | 4.28<br>52.7 | 1.78, 1.64<br>31.3 | 1.28<br>21.8 | 1.58<br>26.8 | 2.91<br>39.6 |
| 21 | R | 8.44 | 4.07<br>55.1 | 1.73<br>27.5 | 1.65<br>24.3 | 3.14<br>40.8 | eNH: NA |
| 22 | C | 8.20 | 4.40<br>58.7 | 3.03, 2.81<br>32.3 | | | NOE: NH-3.64 |

Serines converted to Lan are shown as (S) A; Thr residues converted to MeLan are shown as (T) Abu. Pairs of residues that are crosslinked are shown in the same color.

**Table S5**. Primer used in this study

| Template | Primers | Nucleotide sequence (5` to 3`) |
|---|---|---|
| pET-28-His6-TlaA1-TlaA2-TlaM | ENW_tlaA1_g_R | acctgcaggcgcgccgag |
| | ENW_tlaA1_g_F | aacagattggtggatcggatcctATGAGCCGTAACTATT |
| pET- His6-SUMO | ENW_tlaA1_v_F | gctcggcgcgcctgcaggtcgac |
| | ENW_tlaA1_v_R | ACGGCTCATaggatccgatccaccaatctgttctctgtgagcc |
| pET-28-His6-TlaA1-TlaA2-TlaM | ENW_ tlaA2_g_R | caggcgcgccgagctcgaattcttaA |
| | ENW_ tlaA2_g_F | gctcacagagaacagattggtggatcggatcct |
| pET- His6-SUMO | ENW_pET_ tlaA2_v_F | ctcggcgcgcctgcaggtc |
| | ENW_pET_ tlaA2_v_R | tccgatccaccaatctgttctctgtgagcctc |
| pET-28-His6-TlaA1-TlaA2-TlaM | ENW_ tlaM_g_F | TCACCACATGAATACGAATTTCCGCACTCAGCTG |
| | ENW_ tlaM_g_R | CGACTTAAGCATTATGCGGCCGCAAGCTT |
| pRSF- His6 | ENW_pRSF-v-tlaM-F | TTGCGGCCGCATAATGCTTAAGTCGAACAG |
| | ENW_pRSF-v-tlaM-R | ATTCGTATTCATGTGGTGATGATGGTGATGGCTGC |

**Table S6.** Codon-optimized genes used in this study

| | |
|---|---|
| tlaA1 | atgagccgtaactatttcaaggaggagattaaatactttgctgtcggtcagcgtgatgtcgaagtcagccgcgaagagcttgagcgtgtctcaggcggtaacggttctacagatgactttacctccttagtttgtgattgggcacgcgcaactatcaacgagaattgtgggatgaaaagtagccttgctacgcccctgcaagcgttgt |
| tlaA2 | atgagccgccaagacgccaagaaaactatccaaatgtatgacccggcaggccagaaaaatgtggaaatcagccgtgttgagctggaacgtgtaagcggtggtaacggtgcaaacgaggagatcacaaccctgattaccaccatggtgtgtaaagctattgaggtgacggtgaatgacgggtgcggtatgaagagttctttggccactccatgtaaacgctgt |

| | |
|---|---|
| tlaA2E20D | atgagccgccaagacgccaagaaaactatccaaatgtatgacccggcaggccagaaaaat<br>gtggaaatcagccgtgttgagctggaacgtgtaagcggtggtaacggtgcaaacgaggagat<br>cacaaccctgattaccaccatggtgtgtaaagctattgacgtgacggtgaatgacgggtgcggt<br>atgaagagttctttggccactccatgtaaacgctgt |
| tlaA2E6D | atgagccgccaagacgccaagaaaactatccaaatgtatgacccggcaggccagaaaaat<br>gtggaaatcagccgtgttgagctggaacgtgtaagcggtggtaacggtgcaaacgaggacat<br>cacaaccctgattaccaccatggtgtgtaaagctattgaggtgacggtgaatgacgggtgcggt<br>atgaagagttctttggccactccatgtaaacgctgt |
| tlaM | atgggcagcagccatcaccatcatcaccacatgaatacgaattttcgcactcagctttatcgttc<br>gcttacccttaaagaacgtttcgatcacctgcctaacttaggccaaaagaaggttgattcaattg<br>acgctgaaaaagtgattcacgactggcaaaatgtcagtttttttagatgaaaaaacacttgccaa<br>tcgtttgtcagccaccgatcttgagatgcagcgctttaagtcggcactttatgaaatgtcaagcga<br>tcactggatcaacgaaatgaagaccctgcatcactcgaaattcccctggatggattggcttgaa<br>gaagccttacagctgaatcgcgtgacacccatcccagaggacatcgagaagggcttgcagttt<br>acagtacgcccattcgtactgtgggctaagaagcgcttgaccgattattttggtcagatttccgaa<br>gtcgacccatacatccaaatccataccgtgttggatagtatcctgggtaacctggttgacgggct<br>gaacattatcgcagggcgcacattcgtgcttgaattgcatatcgaacgcgagatgggacaattg<br>gagggagatactcctgaagctcgctttcaaagcttcatccagaaaaagattatgaacccagat<br>cacttggagtttatttactcggagtatccgaccttagcacgtttgttgatgatccgcacccaccactt<br>tatggaggcaattacagaggctattacccgttatttgaacgatcgtaagcaaatcttacaggagt<br>tcaacattcaagacaaacccttaaccgcaatcagcgccgggatgggggactcacatcagcgt<br>tgccgcacggttatgcattttcaattcgaatcggagcaggtgatctacaaaccgaaaaatcttac<br>agtttcgaaccacttccaccaggtgttagattggttgaatgggtgtggttttaccccacccttgagc<br>agttacaaggtcttaaacaaaaaccactatgcgtgggaggaagtagtaacgcaaaaaggat<br>gcagcagtacagaagaggtacagcgtttttatacacgctttggagggttgctggccgtcgtatat<br>agcctttacggaattgatttccattacgagaacatgatcgcaaacggcgaaaaccctatcttaat<br>cgacctggagactttgttccataacagttcctctccgaatgtcgcggaggagatgttggcccagg<br>tcaaggccaacgaccgtttagcgaatagcgtgttgaaaaccgccttgttaccattgtttcacttttc<br>agataaggacggtaaaggcatcgacgttagcggactgggaggtcgcgagcaagagtatccc<br>acaccaattttgcaggtcgaagaatatggaccgaccagatgcgctatgttcgtaaaaatgcg<br>attcttcgtttgagcgggaacctgccccgtttgcacgatcaattgattgacatcaaaccgtatgtcg<br>aatacatcgtcagtggcttcaaacaggcctgccaatttatccaagagcatcaagtgaaattatta<br>tcagatgaaggtcccattgccagttttaagcaagaccaggtccgtatcgttctgcgcaacaccc<br>agttttacgccgacttcctgttggaaacccaacatccggactacttagaagattctctggagcgc<br>gaaaagttgttagatcgcttgtggttcacacaaatgcatgacgaagcgattccatatgagatcg<br>aggatttgttagaaggggatattcccttgtttacggccatgattgacgatacggacctgtacagca<br>gcacggggaagattattccgaactttttcaaagaaagtagttaccagcgcgttatccgccgcatt<br>aagtccttgactcccgatgagattgaacgtcaagcatcctatattacggcatccatcttgggagg<br>aattgagtcgaagactcatctgcaaatcaaacagtatgactttacaccagatcccattaaacat<br>acggaccttccagtccaatctttcgtcgaggaagcagagaagatcggttcgtacttgtccaagc<br>gcgccatctacggagataagaatgacgtgacgtggattggactggccccgaccgcaaataat<br>ctttggactatcgctcctatggattttggtttgtataatggcgtgtgcgggatggccctgttctatagtt<br>atcttgaccaaatctgtaaaaatcgcgaattcggtaatcttgccaaggctgcattgcaaacagct<br>tgcaatgcagggccgttaatccaggatgcgaatgcgttcgtcggccagagctcgatcttgtata<br>cactttctcacatgacggcgttatatggcgaaaaggaagagtggatgtcttccatgaaggagctt<br>cttccaaacattgagcaaaagatcgaacaagatcagcacttcgacttaatctacggaggggc<br>ggggattattcatgtacttcttaacattgctgaacaattcaattgggaacatccattacttatcgcac<br>aaaaggtaggcaaccacttaattaaacacgctatccaaaccgacaacggggtagcatggca<br>tacaggcaaggacaaggcactgttgggcggcttttcgcatgggacttcagggatcgcctggag<br>ccttctgcgtctggcgaacgtttccgggcacgataagtaccacgagtggggtttgaaagccttgc |

| | |
|---|---|
| | agtacgaccgttctctgtatgacgagtcaacgaaaaactggcgtgacattcgtcatgagaagg<br>gcagtagcagtccagtgcagtggtgccacggcgcccctggagtcggcttaggccgtgtgttatg<br>tttaccctatttgcaagaggatccctacatcatcgacgagattagtacctcggtggaaacgacttc<br>aaaagagggaattggtttcagccactcgttgtgccatggtgatctggggaacgcggacttactgt<br>taatggcaggaatcagtttaaagcgtgaggactggatccaaagtgcgcagagtatcgggcat<br>aacgtaattcaaactaagaaaaaacacgggaagtatctgactggcgtatctcactttttggaga<br>ctcccagtctttttctgggattgagcggaattggttatcaacttctgcgtcttgcgtatccagaccaa<br>gtgccgtctgttgtctcgttgcaacctccttttatgaaa |

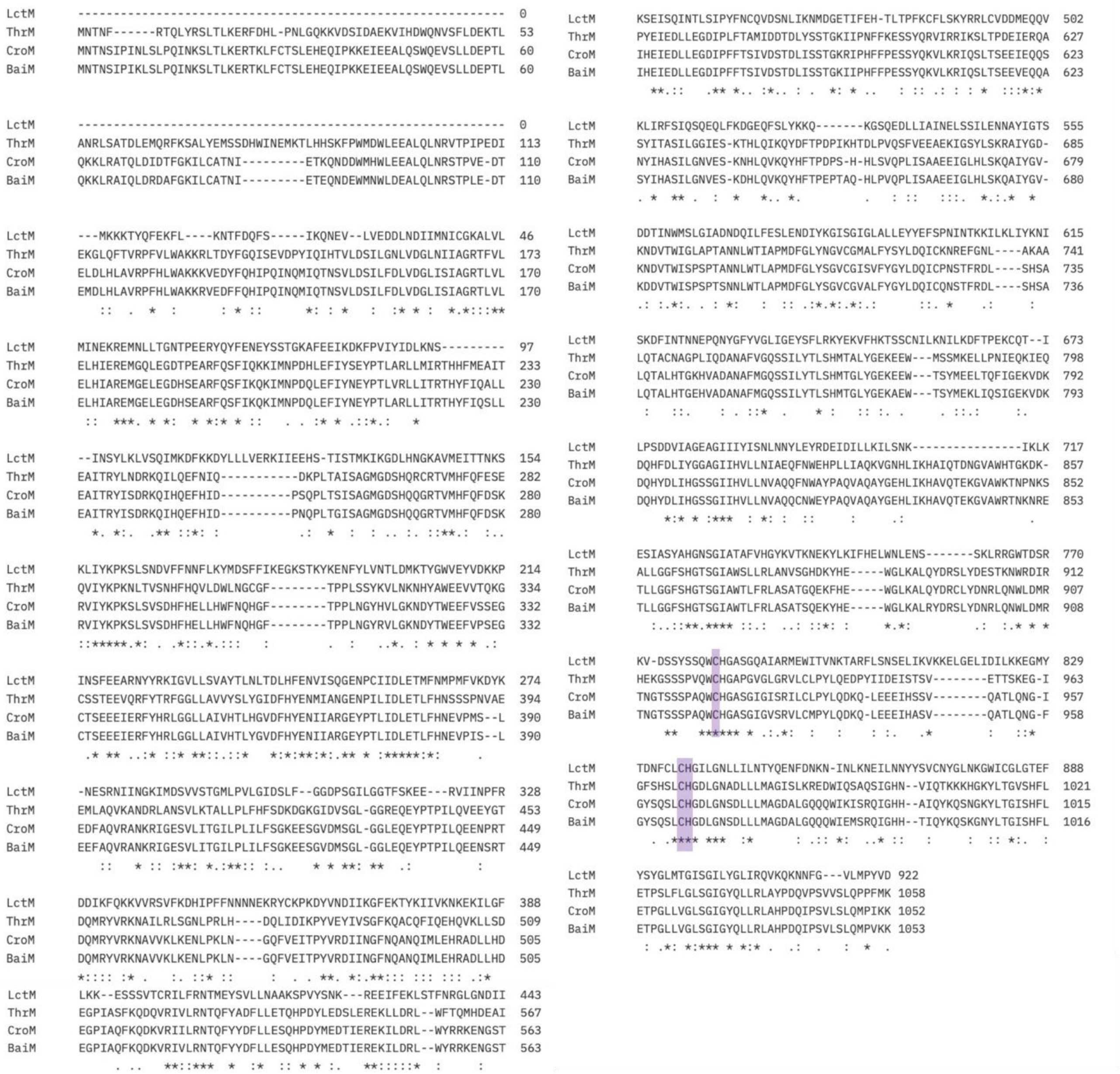

**Figure S1**. Sequence alignment of lanthionine synthases LctM, TlaM, and closely related homologs CroM and BaiM. Highlighted in purple is the Cys-Cys-His triad for the zinc ion binding active site.[1]

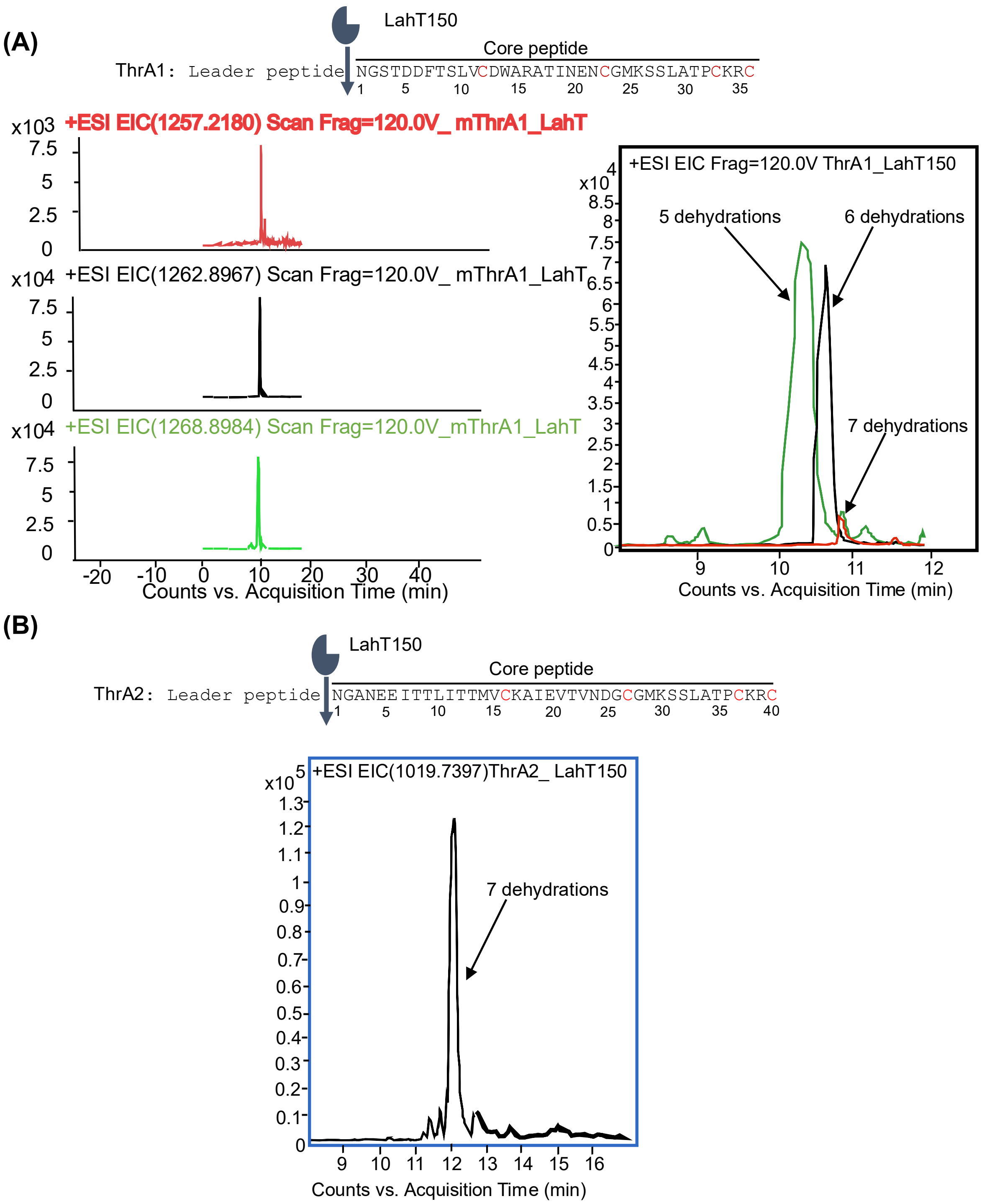


**Figure S2**. Extracted ion chromatograms (EICs) and dehydration pattern analysis by ESI–MS of mTlaAs. **A.** EICs of mTlaA1 and its sequential dehydration products corresponding to losses of water ($[M+H-5H_2O]^{3+}$, $[M+H-6H_2O]^{3+}$, $[M+H-7H_2O]^{3+}$). The chromatographic profiles demonstrate co-elution of dehydration species, with various ratios for five, six, and seven dehydrations. **B.** EIC of mTlaA2 ($[M+H-7H_2O]^{4+}$), illustrating the dominant formation of the seven-fold dehydrated product.

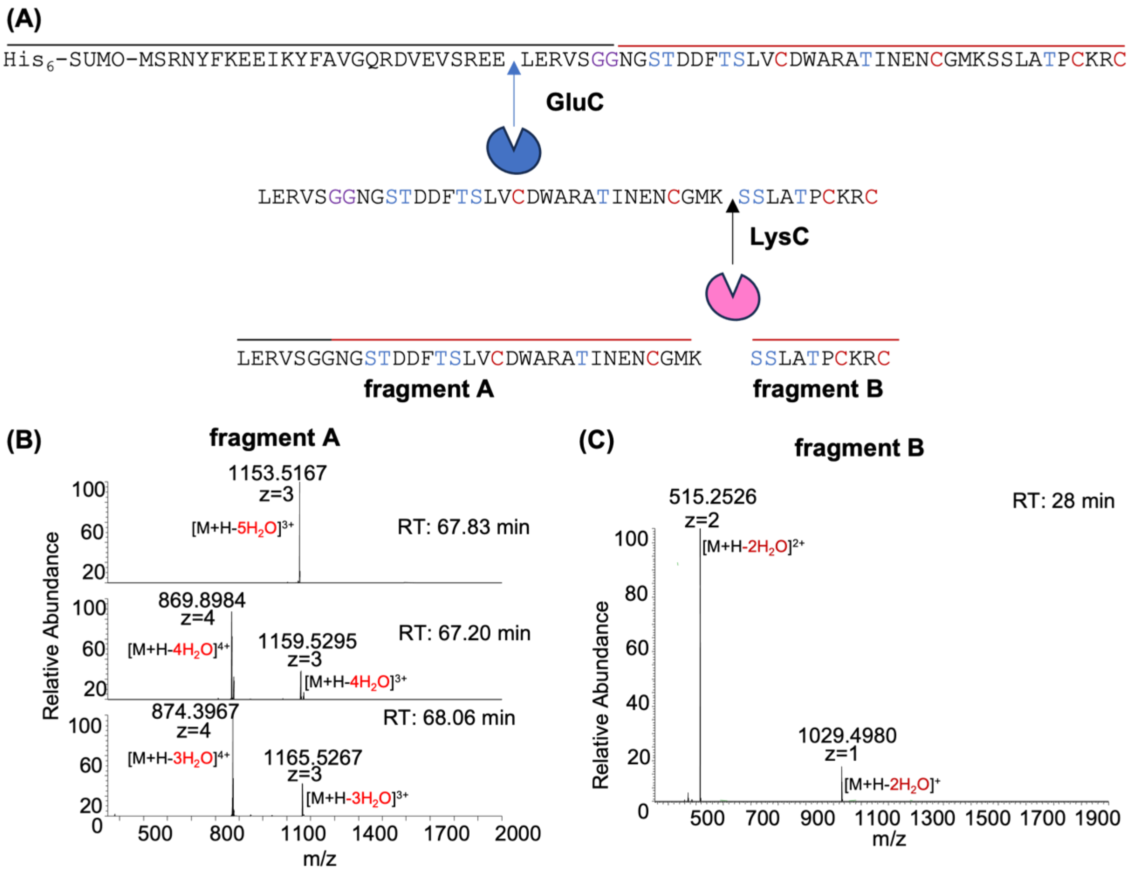


**Figure S3**. **A.** The two fragments from the digestion with GluC followed by LysC of modified TlaA1. **B.** Fragment A contains a mixture of peptides having undergone 3-5 dehydrations. **C.** Fragment B, the C-terminal LysC product of modified TlaA1, contains two dehydrated residues.

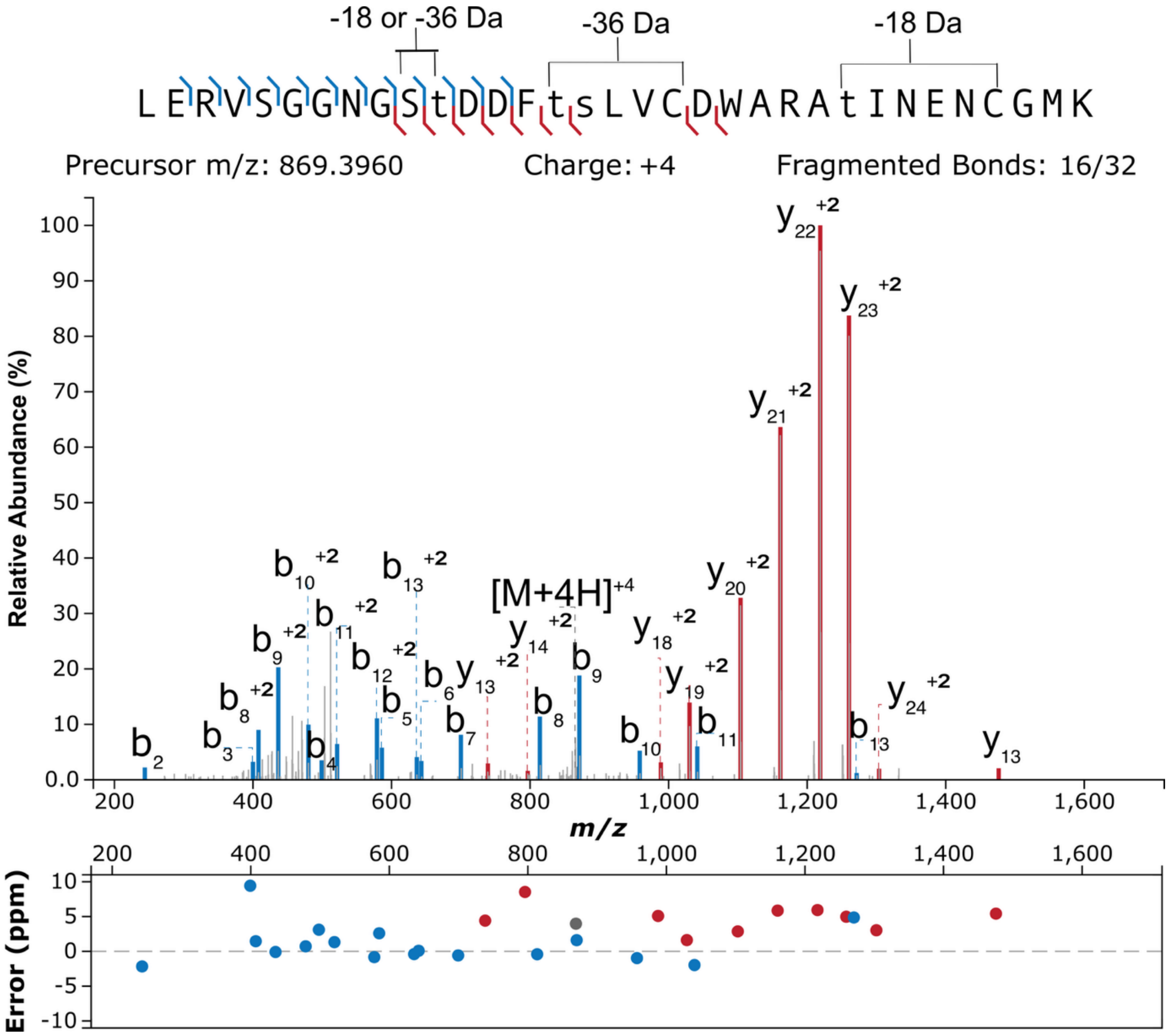


**Figure S4**. Tandem MS of fragment A obtained by GluC/ LysC digestion of modified TlaA1. The four-fold dehydrated peptide was used for fragmentation (corresponding to the six-fold dehydrated full length TlaA1 peptide). Ser3 has escaped dehydration in this peptide. Fragment ion annotation was performed using the interactive peptide spectral annotator[2] with residues indicated in lower case t and s entered as dehydrated.

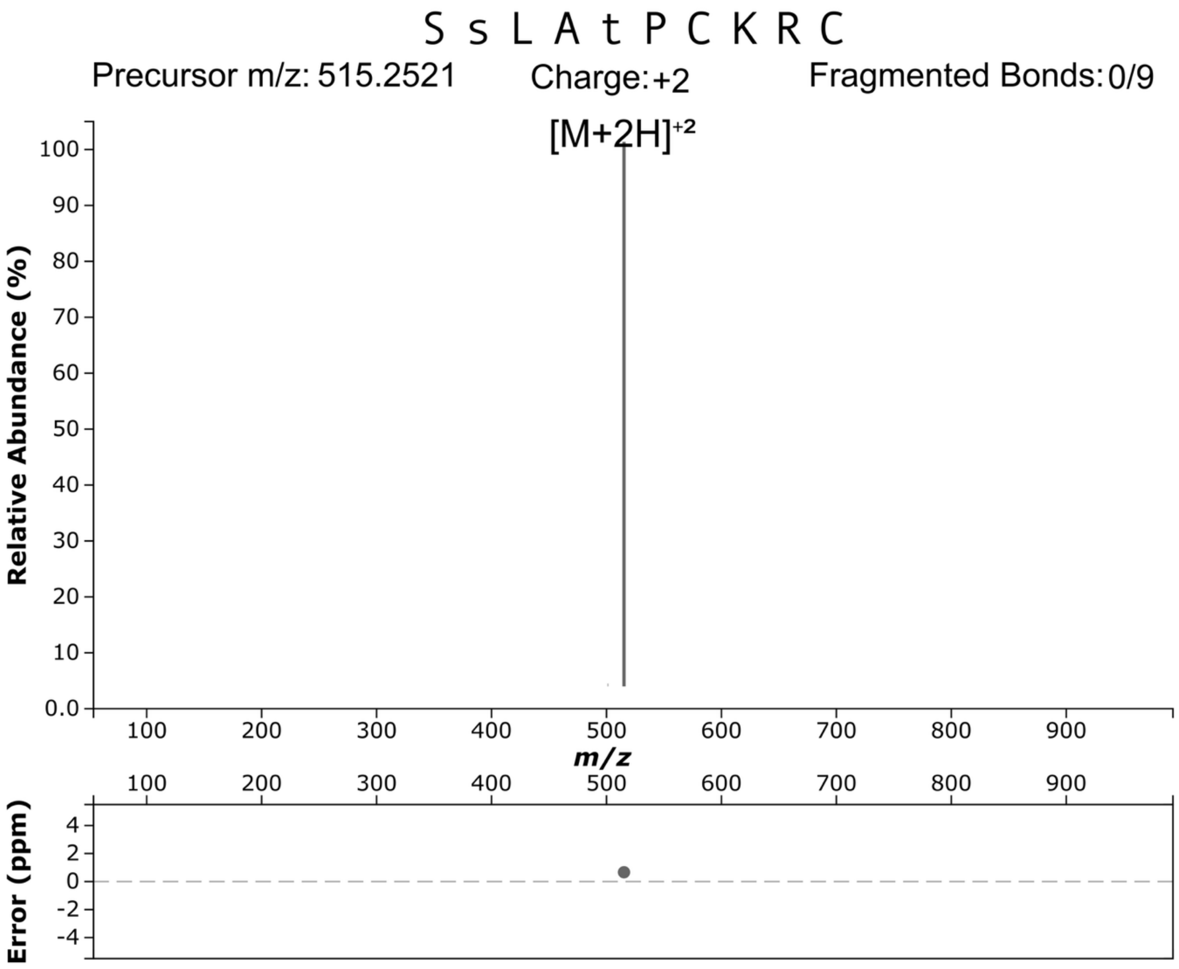


**Figure S5**. Tandem MS of fragment **B** from GluC/LysC digestion of mTlaA1. No fragmentation was observed. The lowercase letters indicate where potential dehydration occurred based on NMR analysis.

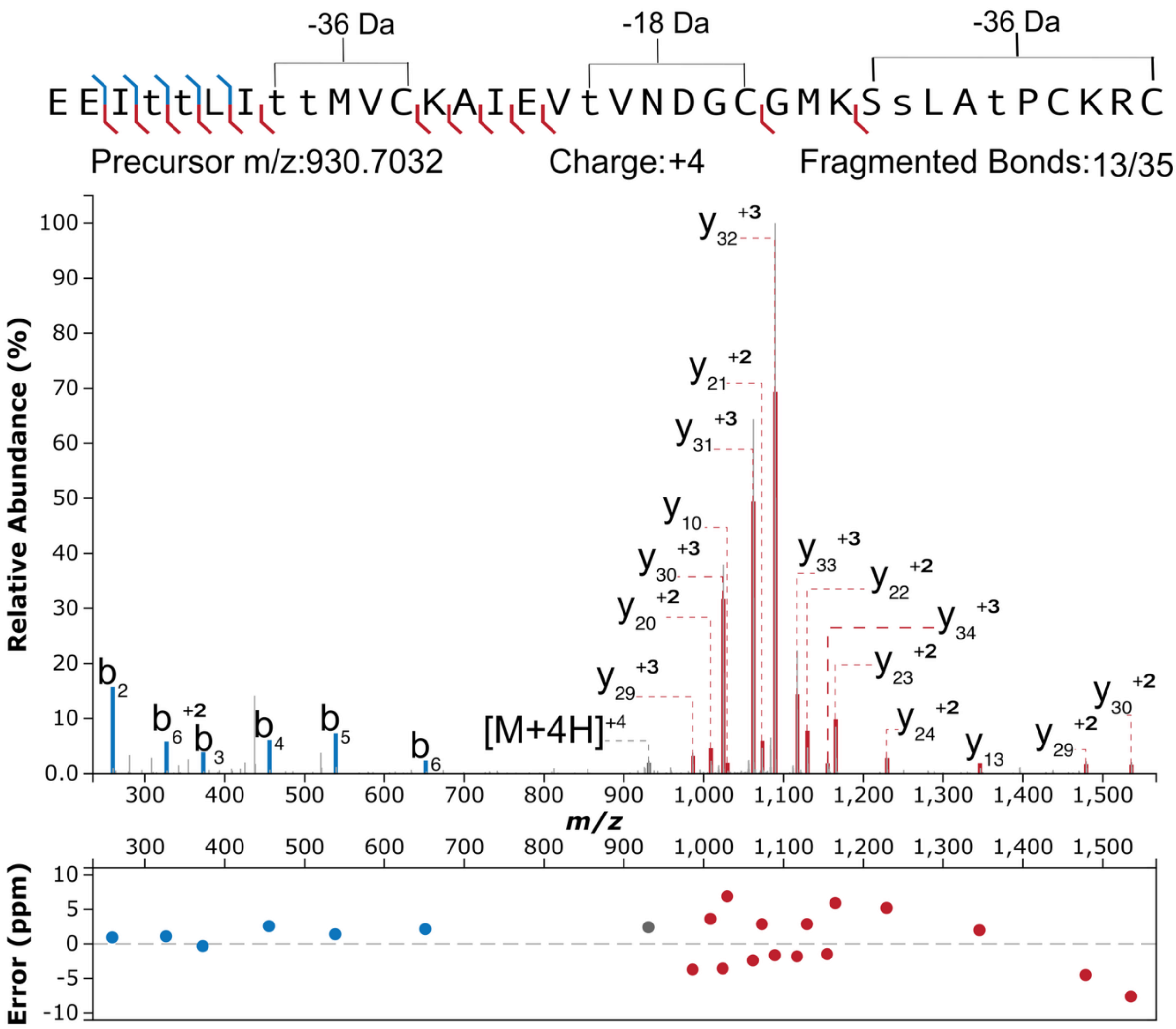


**Figure S6**. Tandem MS of AspN-digested mTlaA2 showing where the post-translational modification occurs. Fragment ion annotation was performed using the interactive peptide spectral annotator[2] with residues indicated in lower case s and t entered as dehydrated. The assignment of dehydration of the Ser and Thr in the C-terminal segment is based on NMR analysis.

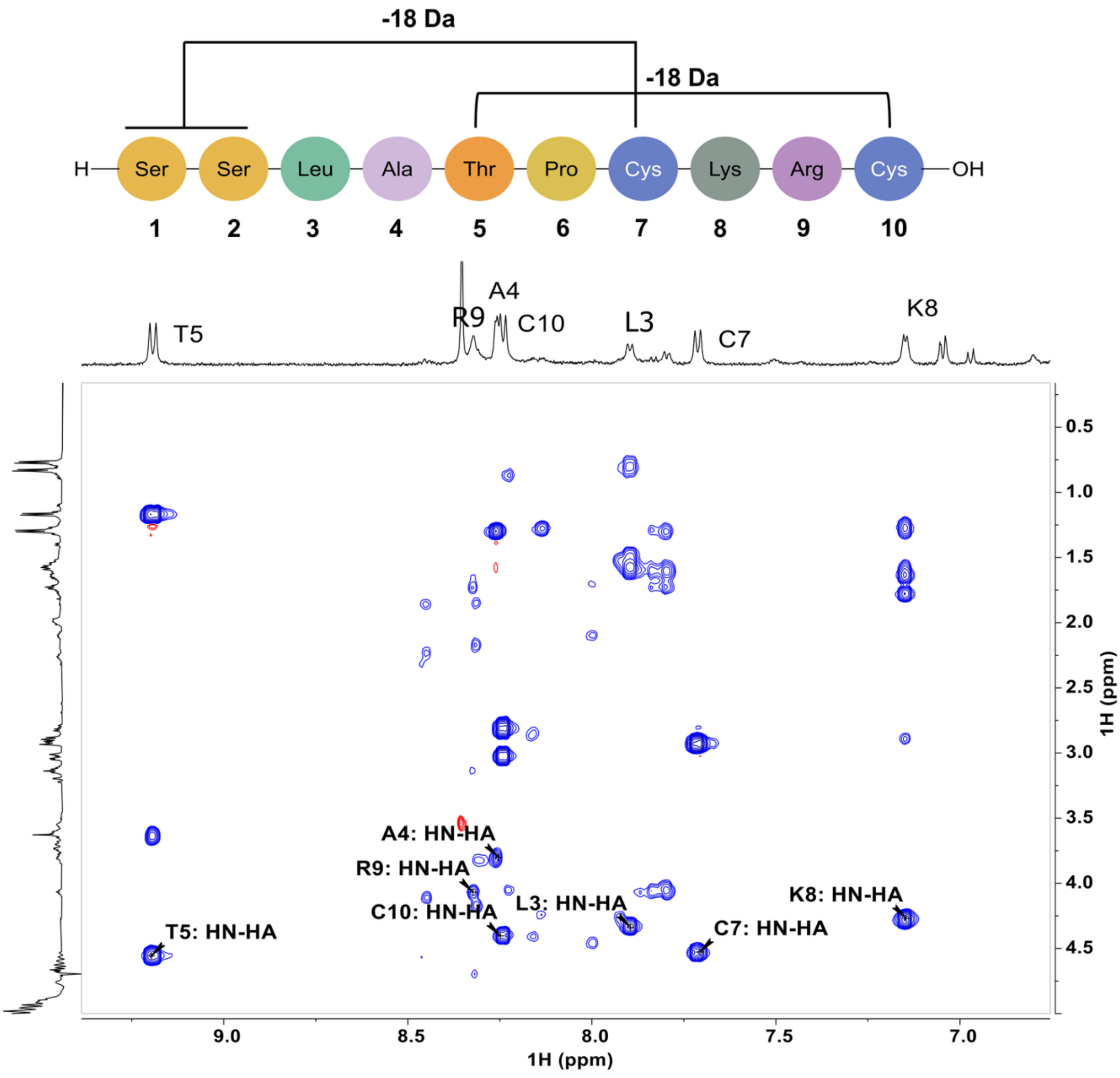


**Figure S7.** $^{1}H$-$^{1}H$ TOCSY spectrum of the 10-residue peptide, fragment **5**. Cross-peaks between the amide and α-protons of each residue are annotated in the figure. The amide proton of Ala2 (formerly Ser2) is not observed.

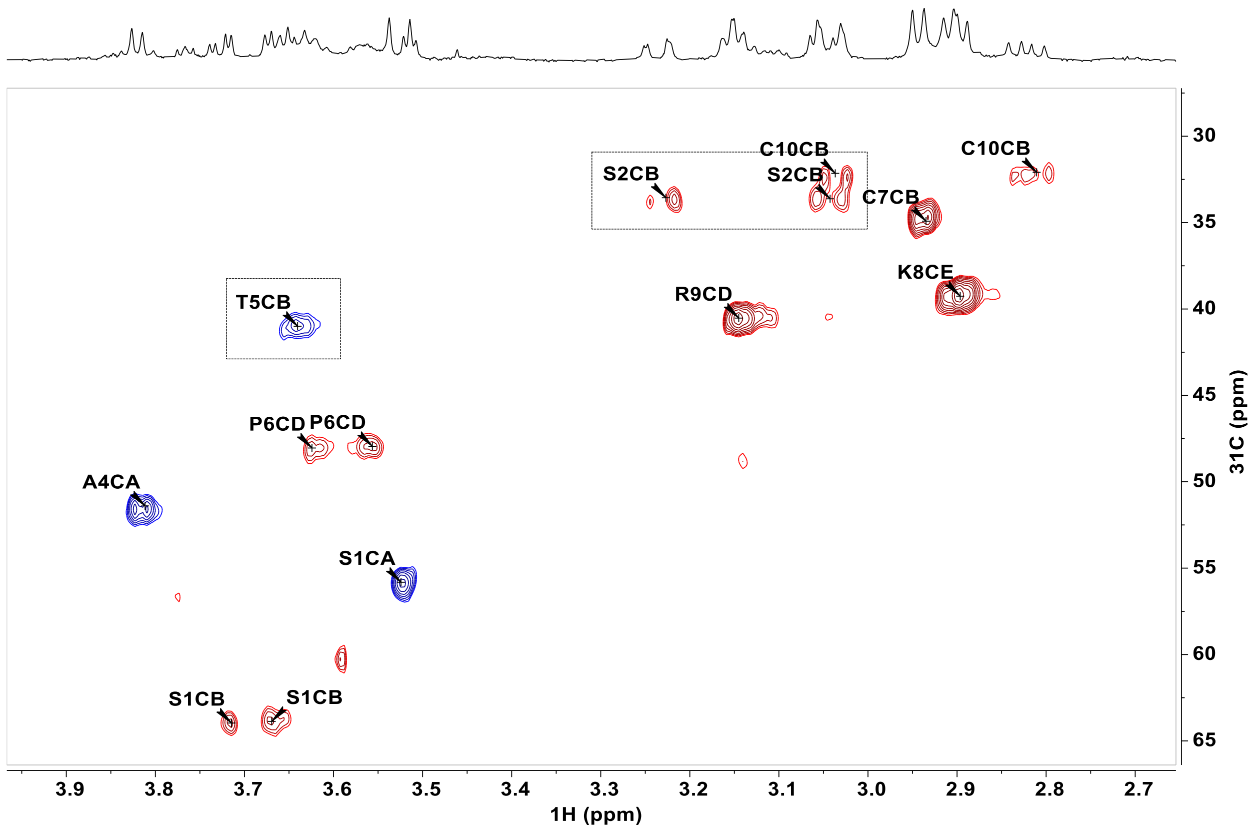


**Figure S8.** ${}^{1}H$-${}^{13}C$ HSQC spectrum of fragment **5** recorded in 100% $D_2O$. Cross-peaks enclosed in dotted brackets exhibit significant deviations in both ${}^{1}H$ and ${}^{13}C$ chemical shifts from the typical values of Ser and Thr residues, indicating their involvement in lanthionine and methyllanthionine formation.

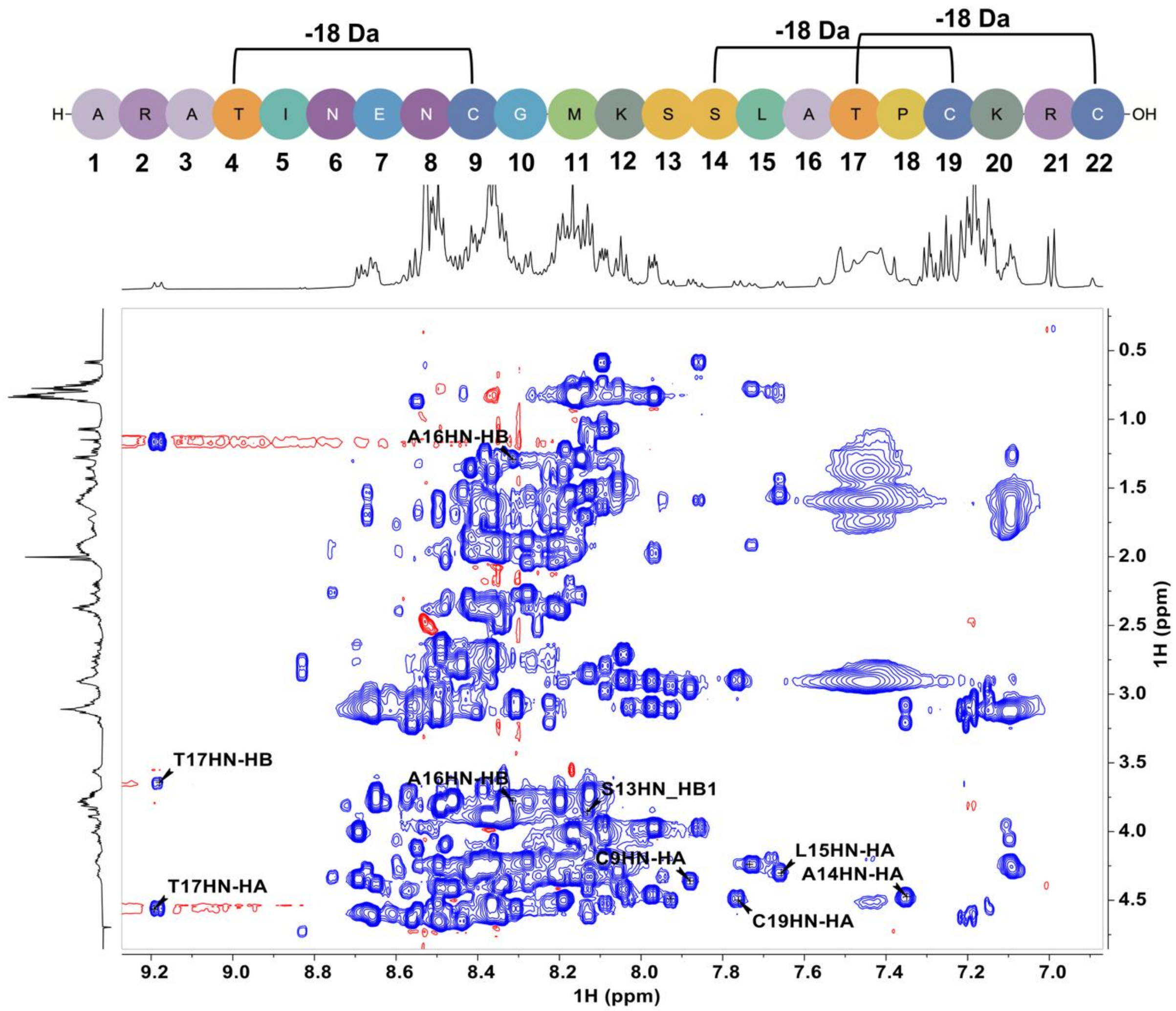


**Figure S9**. $^1$H-$^1$H TOCSY spectrum of the 22-residue peptide, fragment **2.** Cross-peaks between the amide and α-protons of several key residues are annotated in the figure. The residue Ala14 corresponds to the former Ser2 in fragment **5**.

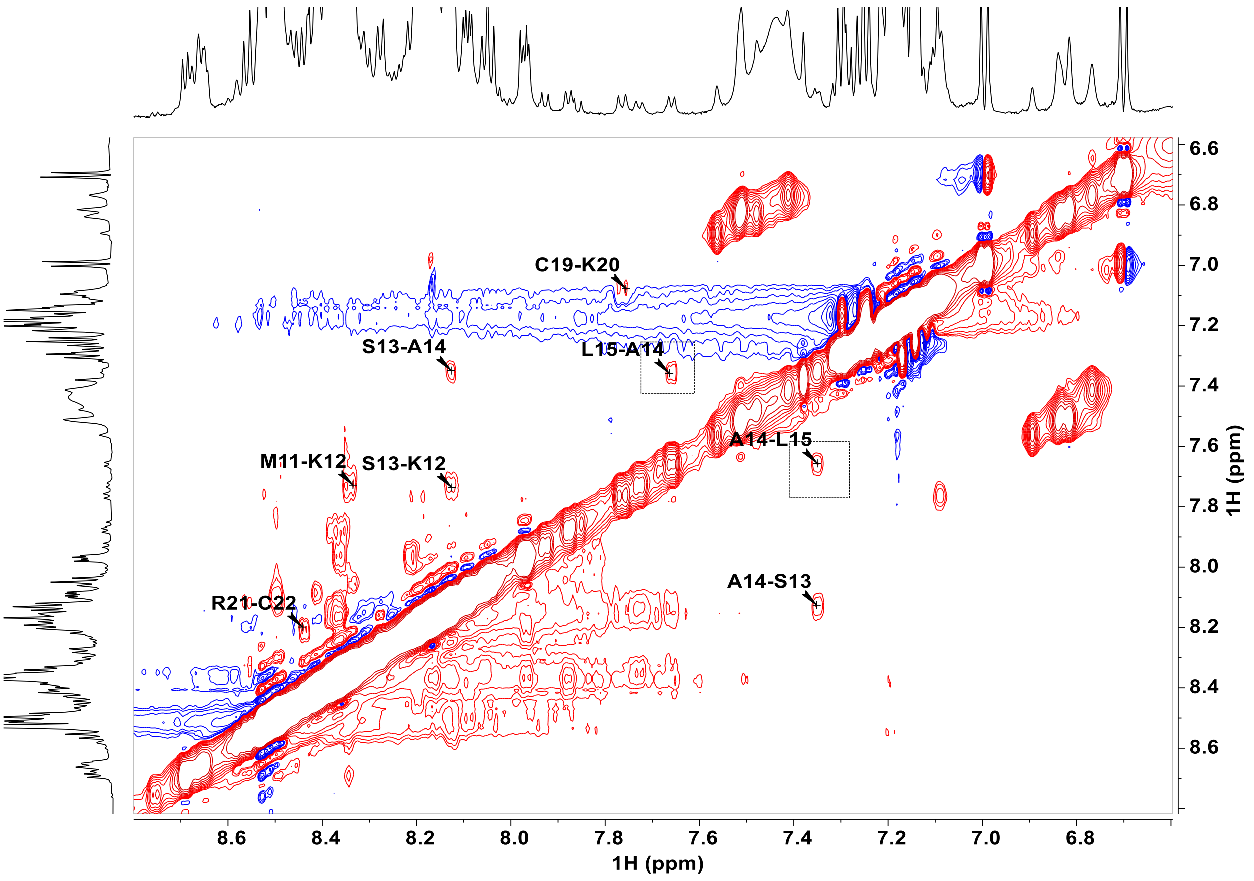


**Figure S10**: The amide region of the $^1H$-$^1H$ NOESY spectrum of fragment **2.** Cross-peaks between the amide protons of Ala14 and Leu15 are displayed in the dotted brackets. The residue Ala14 corresponds to the former Ser2 in fragment **5**.

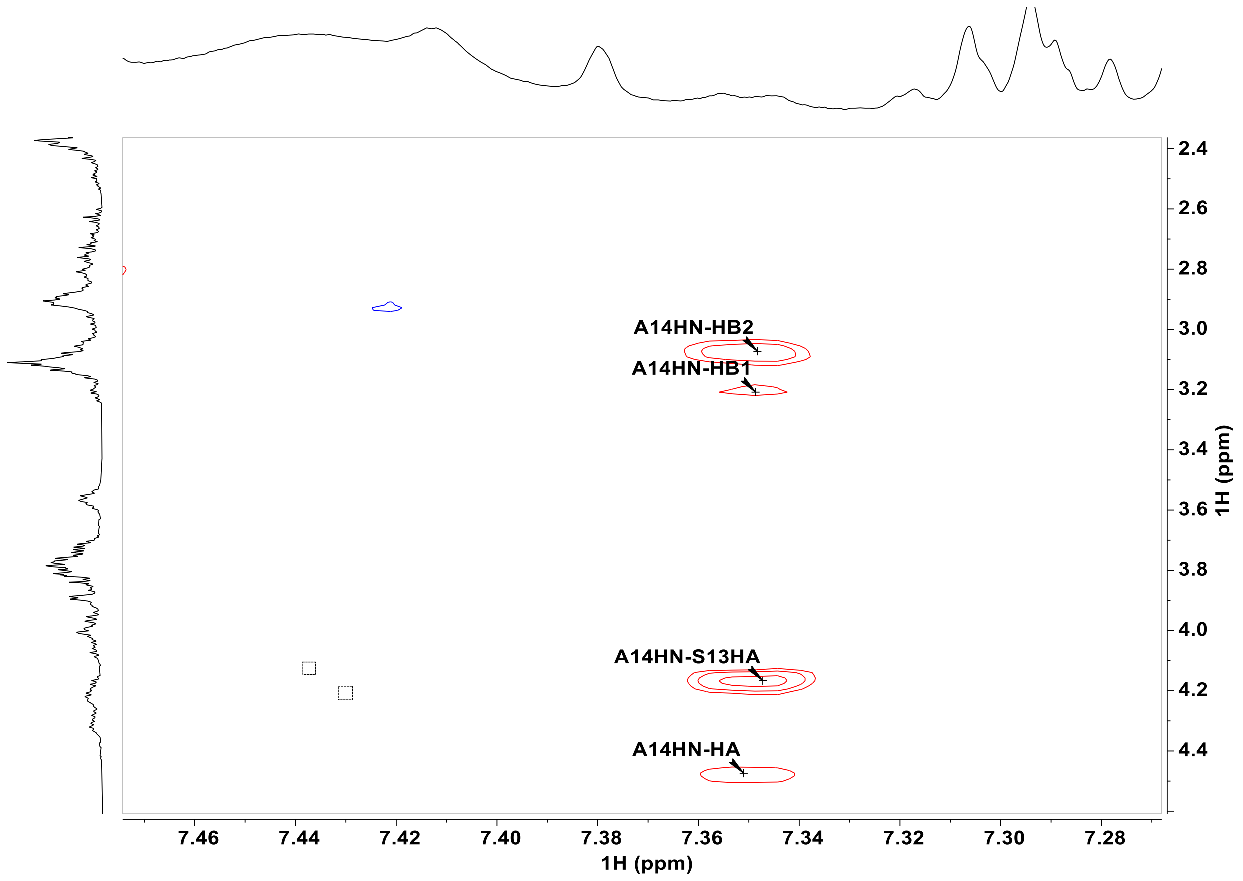


**Figure S11**: The amide region of the $^{1}H$-$^{1}H$ NOESY spectrum of fragment **2**. Cross-peak between the amide proton of Ala14 and a-proton of Ser13 is clearly observed. The residue Ala14 corresponds to the former Ser2 in fragment **5**.

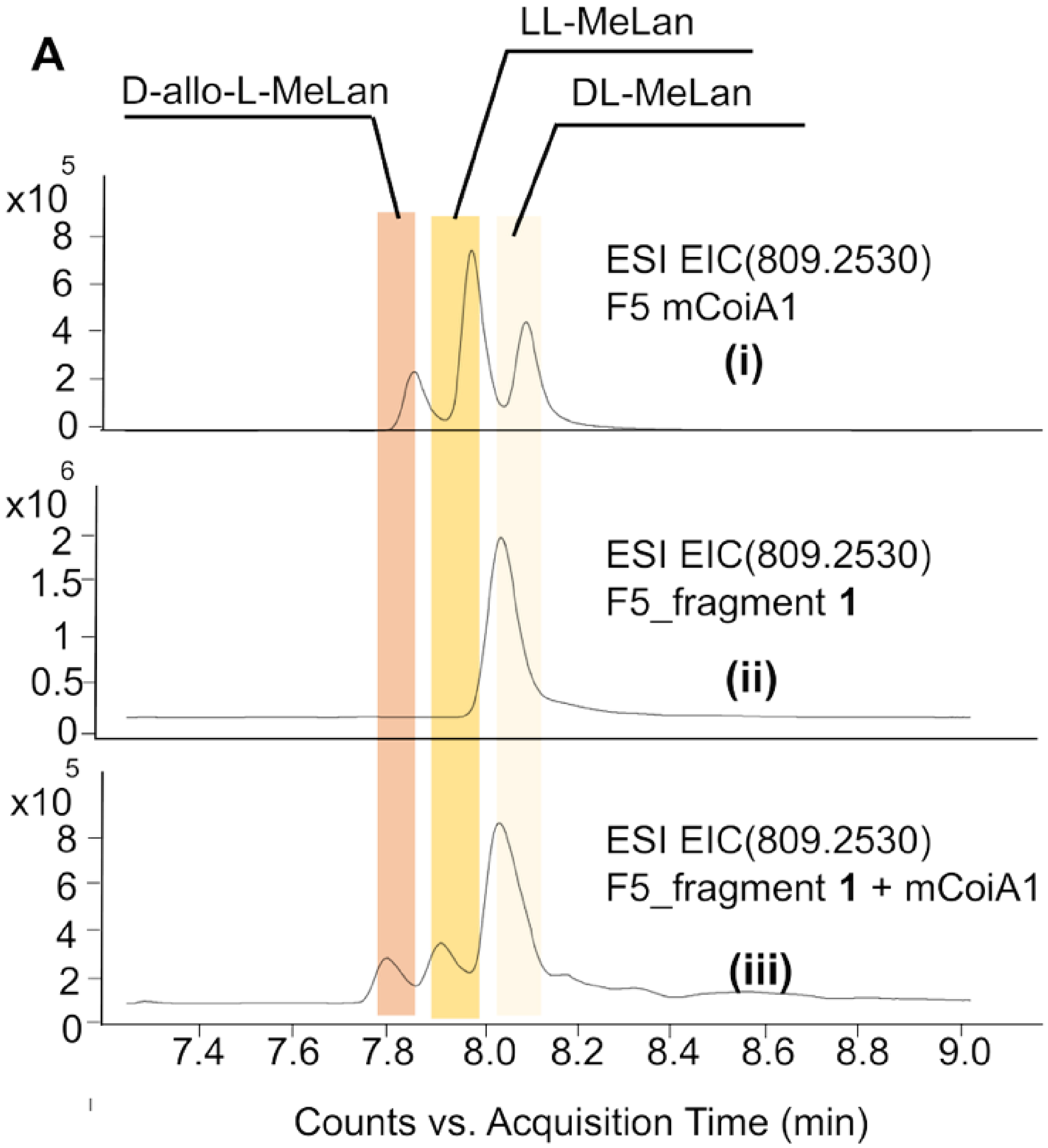
A
D-allo-L-MeLan
LL-MeLan
DL-MeLan
ESI EIC(809.2530)
F5 mCoiA1
(i)
ESI EIC(809.2530)
F5_fragment 1
(ii)
ESI EIC(809.2530)
F5_fragment 1 + mCoiA1
(iii)
x10 5
x10 6
x10 5
7.4 7.6 7.8 8.0 8.2 8.4 8.6 8.8 9.0
Counts vs. Acquisition Time (min)

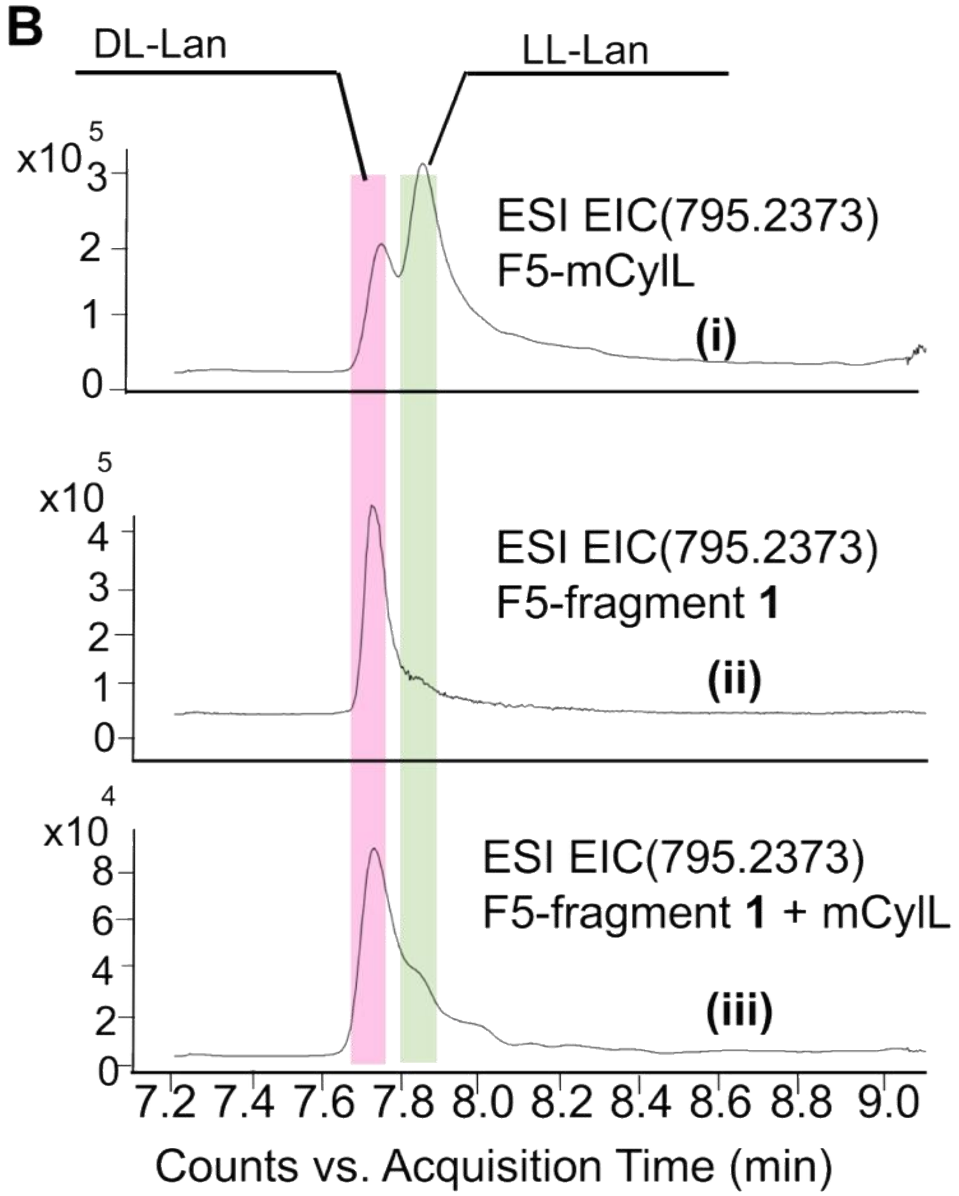


**Figure S12. Marfey's analysis shows that ring A in fragment 1 has the DL configuration.[3] (A)** Marfey's analysis of fragment **1** shows DL-methyllanthionine. mCoiA1 was used as standard.[3] **(B)** Marfey's analysis of fragment **1** also shows the presence of DL-lanthionine. $mCylL_L$ was used as standard.[3] (i) MeLan or Lan standard, (ii) fragment **1**, and (iii) coinjection (fragment **1** + standard).

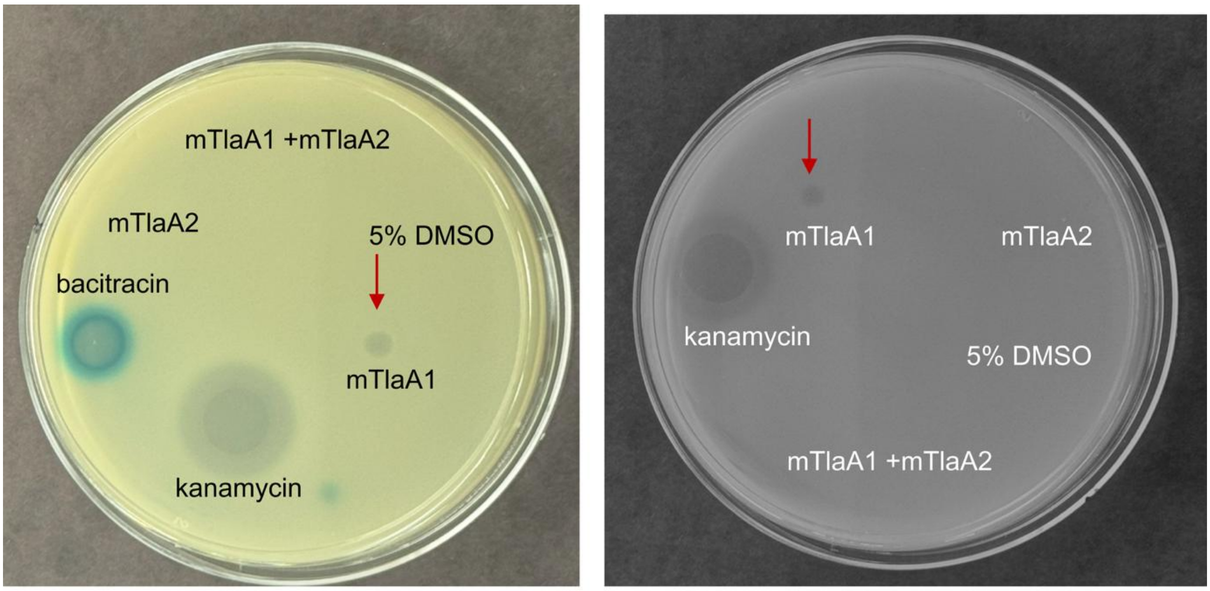


*B.subtilis 2470* *E.coli*

**Figure S13.** Bioactivity screen using agar diffusion assay. (Left). LiaRS assay of mTlaA1 and mTlaA2 cleaved with LahT150 against *B. subtilis* 2470 to observe if the modified peptide targets the lipid II cycle.[4] Since no blue zone is observed, lipid II is unlikely to be targeted. The following samples were spotted: 35 mM bacitracin (positive control, 1.5 µL), 1 mM of mTlaA1 and mTlaA2 cleaved with LahT (2 µL), the negative control 5% DMSO (2 µL), and 4.29 mM kanamycin (1.5 µL). (Right) Bioactivity test against *E. coli*; the same amounts were spotted as in the assay on the left, except bacitracin was not spotted. The activity observed with LahT150-cleaved mTlaA1 is indicated with a red arrow.
